# Pathogen clonal expansion underlies multiorgan dissemination and organ-specific outcomes during systemic infection

**DOI:** 10.1101/2021.05.17.444473

**Authors:** Karthik Hullahalli, Matthew K. Waldor

## Abstract

The dissemination of pathogens through blood and their establishment within organs lead to severe clinical outcomes. However, the within-host dynamics that underly pathogen spread to and clearance from systemic organs remain largely uncharacterized. Here, we investigate the population dynamics of extraintestinal pathogenic *E. coli*, a common cause of bacteremia, during systemic infection. We show that while bacteria are largely cleared by most organs, organ-specific clearance failures are pervasive and result from dramatic expansions of clones representing less than 0.0001% of the inoculum. Clonal expansion underlies the variability in bacterial burden between animals, and stochastic dissemination of clones profoundly alters the pathogen population structure within organs. Despite variable pathogen expansion events, host bottlenecks are consistent yet highly sensitive to infection variables, including inoculum size and macrophage depletion. Finally, we identify organ-specific bacterial genetic factors that distinguish between establishment of within-organ pathogen populations and subsequent survival or expansion.

## Introduction

Sustained bacterial survival in the bloodstream and establishment within otherwise sterile organs is often fatal and severely burdens healthcare infrastructure (1). In healthy individuals, bacteria frequently breach epithelial barriers and enter into the circulatory system, but are rapidly eliminated by complement and other humoral factors and/or captured and killed by liver- and spleen-resident phagocytic cells, clearing the infection and preventing sustained bacteremia (2, 3). However, some pathogenic bacteria can at least temporarily evade or resist these host defenses to potentially enable subsequent dissemination (4–9). Extraintestinal Pathogenic *Escherichia coli* (ExPEC), a set of pathogenic *E. coli* isolates that cause disease outside of the intestine, are leading causes of bacteremia in humans (10). In mouse models of systemic infection, ExPEC strains exhibit markedly delayed clearance from the liver and spleen compared to commensal *E. coli*, suggesting that despite their eventual clearance, ExPEC possess strategies to persist longer within organs, thereby delaying the resolution of infection (4). However, when and where these clearance delays manifest is not fully understood. More broadly, the pathogen population dynamics of systemic infections is highly complex and not well-characterized, and understanding the interplay between bacterial spread, replication, persistence, and clearance can help inform therapeutic interventions (11).

A variety of immunological and microscopy approaches have been employed to investigate the host factors that regulate bacterial dissemination (7, 9, 12). For example, replication of *Streptococcus pneumoniae* within CD169^+^ splenic macrophages (7) or *Staphylococcus aureus* within GATA6^+^ peritoneal macrophages (12) have been shown to enable systemic dissemination. However, such approaches require infection models with very large bacterial burdens and are not easily amenable to high-throughput or population- and animal-wide analyses. Furthermore, these methodologies require prior understanding of within-host pathogen population dynamics during infection, which then enable hypothesis generation to dissect underlying mechanisms of infection.

Several different bacterial population-level approaches have been employed to study infection dynamics (11, 13–15), including systemic infection in *Pseudomonas aeruginosa, Staphylococcus aureus*, and *Streptococcus pneumoniae* (16–18). Most of these methodologies involve barcoding otherwise identical bacteria and enumerating the abundance of barcodes to follow lineages (15, 19). One approach known as STAMP (Sequence Tag Based Analysis of Microbial Populations) leverages deep sequencing of bacteria barcoded with random DNA sequence tags and population genetics frameworks to quantify bottlenecks, dissemination patterns, and, more recently, replication rates (20, 21). STAMP was initially developed to study *Vibrio cholerae* intestinal colonization, and subsequently applied to *Listeria monocytogenes* dissemination from the intestine, *Pseudomonas aeruginosa* bacteremia, and *Streptococcus pneumoniae* nasopharyngeal colonization (16, 22–24). We recently created STAMPR, a successor to STAMP that relies on a new computational framework and employs additional metrics to study infection population dynamics at high resolution (25).

Previous analyses of infection population dynamics have often been limited either by a small number of barcodes, organ samples, or time points. Here, we leveraged STAMPR to comprehensively map ExPEC population dynamics following dissemination in blood after intravenous (i.v.) inoculation using ~1100 barcodes across over 400 samples from 6 time points and 12 within-animal compartments. Intravenous bacteremia models vary in outcome depending on strain, dose, and length of time post inoculation (4, 17, 26, 27). We adopted a non-lethal dose of ExPEC to investigate the delayed clearance of this pathogen and monitored the clearance phase to uncover the population dynamics that underlie delayed clearance. Although ExPEC is largely cleared by most organs, some organs, most notably the liver, stochastically fail to clear the pathogen. We show that these failures are attributable to the massive expansion of very few bacterial cells from the inoculum and explain the marked variation in bacterial burden observed between animals and organs. In certain instances, these clones disseminate and alter the population structure of distal organs. Despite these dramatic clonal expansion events, the magnitude of host bottlenecks remains consistent within organs between animals. However, bottleneck sizes are plastic and highly sensitive to experimental conditions. Both macrophage depletion and increased dose widen the bottleneck but stochastic clonal expansion events can obscure CFU-based detection of these phenotypes. Finally, guided by our granular assessment of population dynamics, we identify ExPEC genes required for the establishment of and persistence through infection as well as clonal expansion; unexpectedly, bacterial hexose metabolism distinguishes these processes in an organ-specific manner. Collectively, our findings explain the population-level phenotypes that underpin ExPEC systemic dissemination and deepen our understanding of the origins of within-host bacterial populations.

## Results

### Experimental design and analyses

A key metric for understanding within-host population dynamics is the founding population size (FP) (20, 21). FP quantifies the number of unique cells from the inoculum that give rise to the pathogen population at a sampling point. Higher FP values are indicative of wider (more permissive) bottlenecks. FP can be measured by an equation derived by Krimbas and Tsakas (28), the output of which is referred to as N_b_. The greater the difference in barcode frequencies between input and output samples, the smaller the N_b_. In STAMPR, an algorithmic adjustment is made to N_b_ that limits the influence of disproportionately abundant barcodes, resulting in a new metric called N_r_ (25). The degree to which N_b_ is biased by these “outliers” can be exploited by calculation of the N_r_/N_b_ ratio, providing a measure of uniformity of barcode frequencies; this ratio is larger in samples where a few bacterial clones have markedly expanded.

We introduced ~1150 20nt random and unique sequence barcodes into the ExPEC (strain CFT073 rpoS^+^) (29–31) *lacZ* locus. The barcodes were stably maintained in the absence of selection for at least ~50 generations (Figure S1A) and did not modify the growth rate of the strains in LB media compared to the parental strain (Figure S1B). Deep sequencing of the barcodes at different known bottleneck sizes (i.e., plated CFU) revealed that N_r_ and N_b_ both accurately quantified in vitro bottlenecks up to 10^5^ CFU for this library, confirming the fidelity of these metrics and the absence of technical artifacts during library preparation (e.g., PCR jackpots, contaminating barcodes) (Figure S1C).

Mice were injected with 10^7^ CFU of the barcoded ExPEC library and were sacrificed beginning 4 hours post infection (hpi) and then daily up to 5 days post infection (dpi) (Figure 1A). The blood, bile, left and right halves of the spleen, left and right kidney, left and right lung, and all four lobes of the liver were harvested at each time point. 4 mice each day were sacrificed from 0 to 4 days post infection, and 12 mice were sacrificed at 5 dpi, collectively yielding 384 CFU measurements across 12 sub organs and 6 time points. Sequencing for STAMPR calculations were carried out on 322 samples that had non-zero 0 CFU. Organs containing as few as 1 CFU were sequenced and analyzed because such samples served as important within-run sequencing controls, yielding information about the level of noise due to technical artifacts (such as index hopping), as they should only possess one highly dominant barcode.

**Figure 1.**
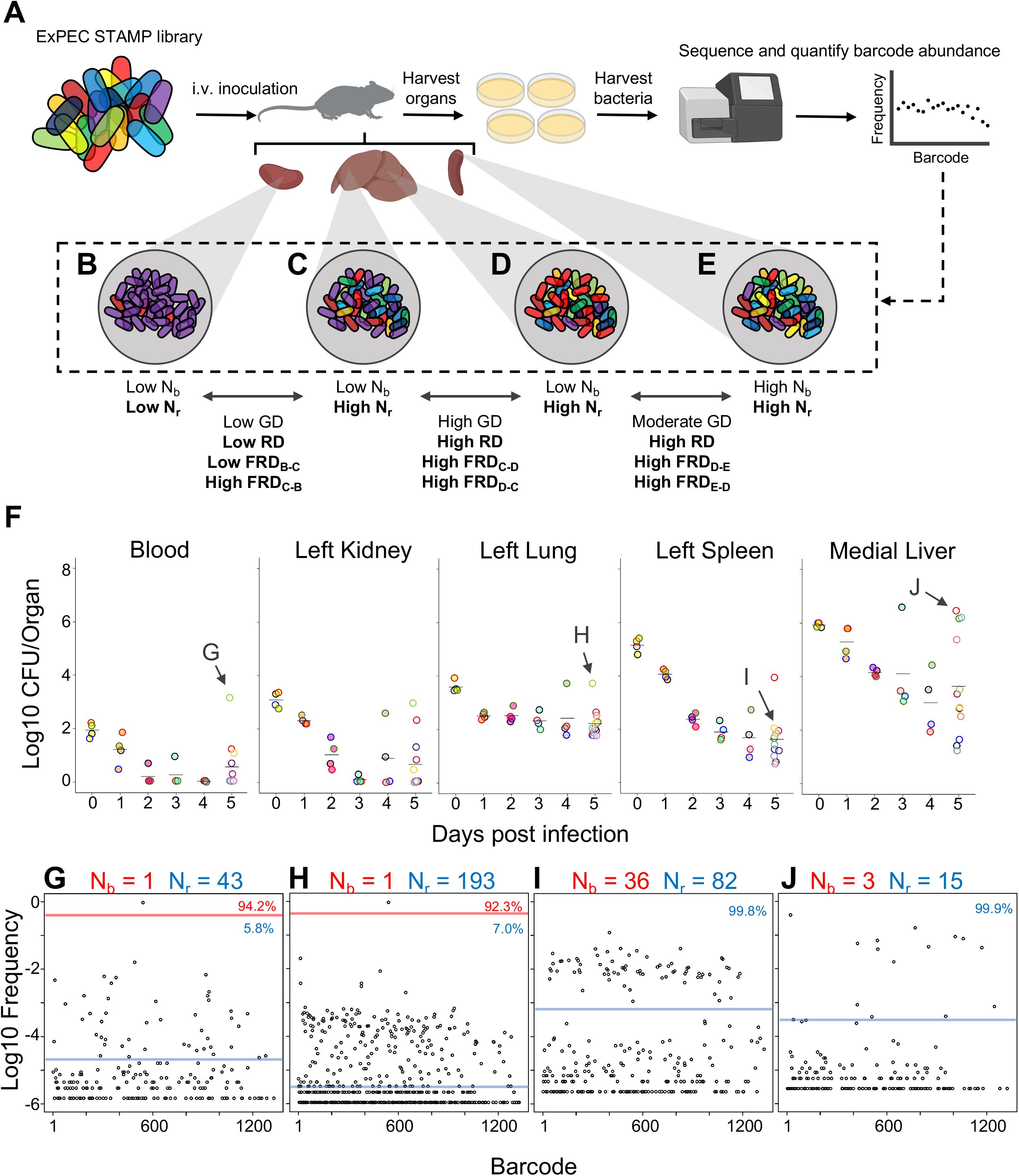
Experimental design and metrics and CFU dynamics. A) The experimental schematic depicts the ExPEC barcoded library as colored bacteria, where colors indicate unique barcodes; the library was injected i.v. into mice, after which organs were harvested, homogenized, and plated for enumeration of CFU. Bacterial cells were scraped, pooled, and the barcode abundances in the output populations were quantified. The distributions of the barcodes (far right graph) define the population structure of the organ, schematized in B-E. B-E) Graphical depiction of populations with different combinations of N_b_ and N_r_ values. These populations arise from B) a tight bottleneck and subsequent expansion of purple cells, C-D) a wider bottleneck and expansion of purple (C) or red (D) cells and E) a wide bottleneck and even growth of all cells. Metrics used for comparisons between samples (GD, RD, and FRD) are indicated. F) CFU recovered from select organs on days 0-5; the full CFU data set is provided in Figures S2-S4. Points with the same fill and border color were obtained from the same animal. Arrows pointing to G-J, correspond to the points where complete barcode frequency distributions are shown below (in G-J). G-J) Barcode distributions and N_b_ and N_r_ values are shown. Red lines separate populations that were distinguished by the N_r_ algorithm (25). The blue line indicates the algorithm-defined threshold for noise. Percentages represent the relative abundance of barcodes within each region. G and H represent examples with highly abundant barcodes that skew N_b_ values lower.

### Bacteria fail to be cleared in a fraction of animals

We first address the patterns of bacterial burden up to 5 dpi across the 12 compartments. As observed in previous studies (4), the highest CFU values were found in blood and most organs 4 hpi (0 dpi) and few bacteria remained detectable in blood after 1dpi (Figure 1F, Figure S2). The kidneys displayed a similar trend, although some bacteria were detected up to 5 dpi in some animals (Figure 1F, Figure S3). The spleen contained more bacteria than the kidneys but still trended towards clearance (Figure 1F, Figure S2). In the lung, however, even at 5 dpi, several hundred bacteria were present, suggesting that the lung has lower clearance capacities compared to the other organs (Figure 1F, Figure S3). The rapid pathogen elimination from blood likely reflects clearance by filtrative organs like the liver and spleen, which had the highest bacterial burdens 4 hpi, rather than killing by blood (e.g. complement), since mouse blood is permissive for E. coli growth (32).

Clearance failures were most prominent in the liver, which yielded the highest CFU values 5 dpi (Figure 1F, Figure S4). Hepatic bacterial burdens were bimodal, with some mice containing ~10^6^ bacterial cells while littermates had ~10^4^-fold fewer bacteria. Notably, animals with high CFU had visible abscesses in the liver. Lobes of the liver that lacked visible abscesses did not have high CFU, arguing against the presence of occult abscesses. While CFU were usually cleared from organs outside the liver, high CFU were also occasionally observed in additional organs; for example, in one animal, the bacterial burden in the left and right lungs differed by 100-fold (Figure S3). Unlike the liver, the organs with elevated CFU did not harbor visibly apparent abscesses. We also observed 3 animals with very high bacterial burdens in the bile (exceeding 10^7^ CFU/ml) aspirated from gallbladder samples from 4 and 5 dpi (Figure S2). Taken together, measurements of bacterial burden alone suggest a model whereby the pathogen distribution early during infection is highly predictable; however, by 5 dpi there is substantial variability in burden, and the range of this variability is organ-specific. To generalize, we can identify at least two trajectories from CFU alone. The first is successful clearance, where very few bacteria survive or persist till 5 dpi with a general decrease in CFU over time. The second appears to be variable organ-specific clearance failure, the most dramatic of which result in liver abscesses but were also detected in “outlier” animals at least once in all organs.

### Clearance failures are mostly driven by clonal expansion

To determine the ExPEC population structure within organs, barcode frequencies were determined and N_b_ and N_r_ were calculated from the 322 samples that had non-zero CFU (Figure 1G-J, Figure S2-S4). Figure 1G-J demonstrates examples of N_b_ and N_r_ values calculated from barcode frequency distributions. The distributions themselves provide intuitive snapshots of the bacterial population structure in single animals. Figure 1G demonstrates an animal with unusually high CFU in blood, where ~95% of reads corresponded to a single tag. The same barcode was highly abundant in the left lung of the same animal (Figure 1H), accounting for more than 90% of reads. In addition, this sample had an underlying population that comprised 7% of reads. Therefore, the elevated bacterial burden in the left lung of this animal results from a clone that was also circulating in blood. Figure 1I depicts bacteria from a spleen sample from a different animal (than Figure 1G) in which ~80 barcodes represented ~100 CFU, revealing that there was no clonal expansion (Low N_r_/N_b_ ratio), similar to Figure 1E. In stark contrast, Figure 1J represents a liver lobe with an abscess (from a different animal than Figure 1I) with >10^6^ CFU derived from only ~10 tags, an example of dramatic replication of very few cells.

The patterns described above (exemplified by Figure 1B-E and 1G-J) can be distilled by comparisons of N_b_, N_r_, and CFU values. For example, when N_b_ and N_r_ are similar to each other and to CFU, we can deduce that the bacterial population has undergone very little replication (e.g., Figure 1I). In contrast, organs with high CFU and low N_r_ and N_b_ values contain populations that resulted from expansion of very few cells (e.g., Figure 1J, and Figure S2,S3,S4). At 4hpi, when mice were still bacteremic, we observed that the number of blood CFU (~100) was nearly the same as the corresponding N_r_ (~80), indicating that at this point most bacteria possessed different tags, suggesting that there has been little replication following inoculation (Figure S2). Like the blood, at early time points both kidneys had N_r_ values similar to CFU, indicating minimal net replication (Figure S3). Our approach also permits analysis of the underlying populations in animals that appear to be “outliers” from others in the cohort. One example is an animal from 3 dpi where the left kidney possessed 0 CFU and the right had >10,000 CFU (Figure S5A). The population in the right kidney was represented by only 8 tags, revealing that ~8 bacterial cells from the inoculum had dramatically expanded (Figure S5B). In the spleen at 4 hpi (0 dpi), N_r_ and N_b_ values were ~10^4^, slightly less than CFU values (Figure S2). By 5 dpi, the CFU recovered (~20) was very close to values of N_r_, indicating that these remaining cells had not undergone substantial replication. One notable exception was a sample from 5 dpi (mouse 23) that had a much higher bacterial burden than other mice (Figure 2A). Since this animal also had a markedly elevated N_r_ (wider bottleneck) as well as an elevated N_r_/N_b_ (expanded clones over a diverse population), the increased bacterial burden can be attributed both to clonal expansion as well as a wider bottleneck, which is consistent with visual observations of barcode frequency distributions (Figure 2BC). In the lungs, CFU approximated N_r_, indicating that the bacterial population had undergone very little replication (Figure S3). This is consistent with the longitudinal CFU measurements, which plateaued by 3 dpi (Figure 1F).

**Figure 2.**
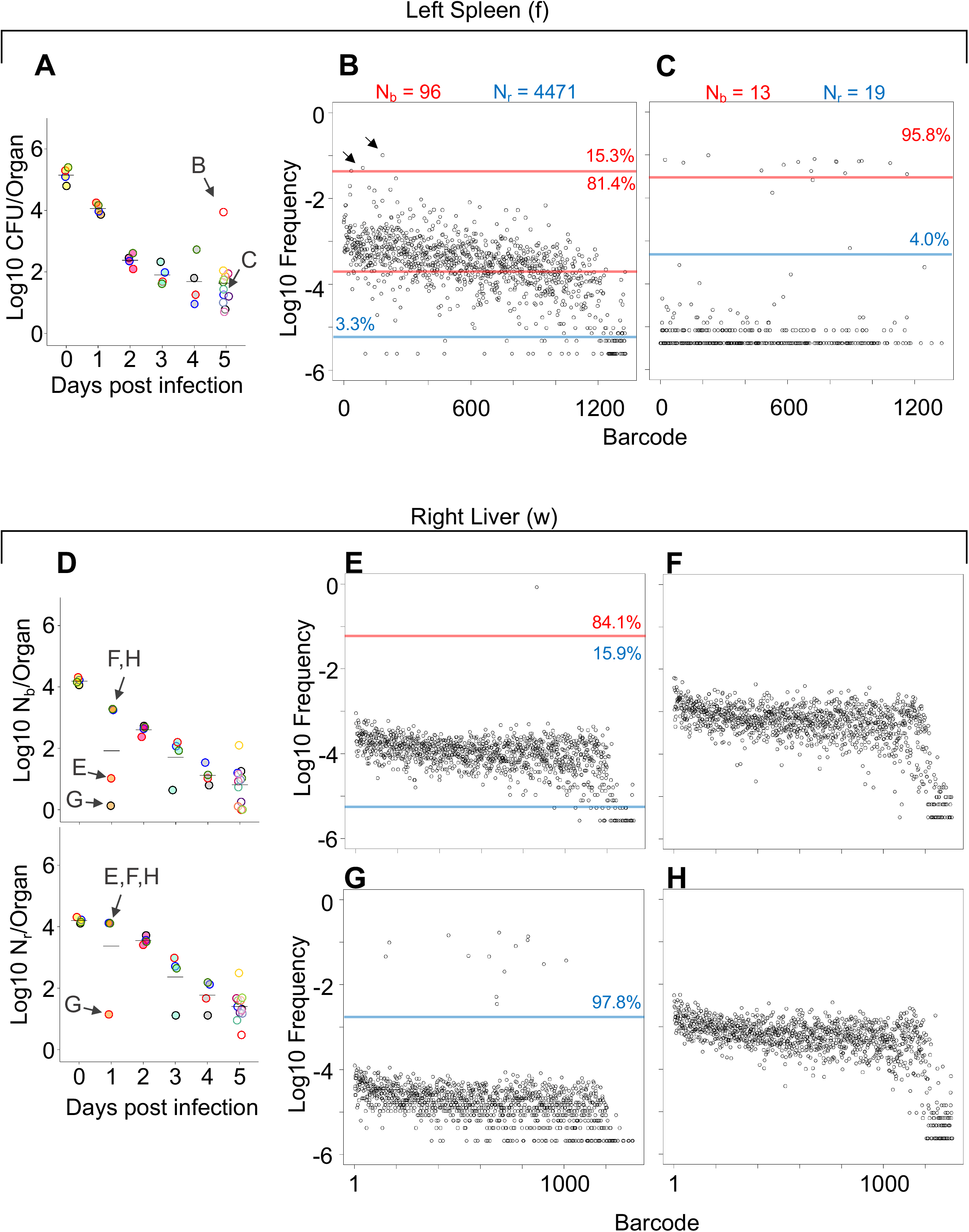
Comparisons of barcode distributions in animals with or without clonal expansion events. A) CFU from the left half of the spleen 5 dpi; points with the same fill and border color are derived from the same animal. B,C) The underlying barcode distribution found in animals B and C are shown in the respective graphs on the right. The highly increased bacterial burden in animal B relative to animal C can be attributed to both a wider bottleneck and clonal expansion. In the barcode frequency distribution shown in B, a diverse population is evident, as is the clear expansion of a few clones (arrows). In contrast, in animal C, N_b_ and N_r_ are similar to CFU, suggesting that the population has undergone relatively little replication. D) N_b_ and N_r_ plots from the right lobe of the liver 1 dpi. The underlying barcode distributions in animals E,F,G, H are shown in the respective graphs to the right. Animals E and G have clonal expansion events; expansion is so substantial in G (~98% of reads) that the algorithm cannot distinguishing the underlying population from noise, and N_r_ is not substantially higher than N_b_ for this sample. In contrast, N_r_ can identify this population in sample E. Red and blue lines denote sub-populations and noise, as in Figure 1.

As discussed above, some of the largest bacterial burdens in these experiments were present in the liver at 5 dpi (Figure 1F, S4). The elevated hepatic CFU values were associated with clonal expansion events, since N_r_ and N_b_ were orders of magnitude lower than CFU in animals with abscesses (Figure 1J, Figure S4). Even at 1 dpi, two animals had clonal expansion (N_r_/N_b_ >1000) in multiple lobes (Figure 2EG and Figure S4) where ~1% of barcodes comprised a large majority of the reads (Figure 2E-H). Nevertheless, at 5 dpi all animals had similar N_r_ values, despite some having abscesses and the concomitant 10,000-fold increase in bacterial burden (Figure S4). Thus, the immune and physical factors that govern bottlenecks appear to be highly consistent between animals. Instead, the focal and stochastic expansion of a few clones within abscesses, rather than a general increase in bacterial survival, explains the wide variance in liver CFUs. In general, these analyses revealed that N_r_ values were more consistent and lower than CFU across all organs particularly at 5 dpi, revealing that most observations of very high bacterial burden are driven by very few bacterial clones (Figure S2, S3, S4). Apart from the lung, FP sizes decreased during the 5 days of observation. Therefore, the host forces that restrict the bacterial population in this model act throughout the infection and are not limited to the initial establishment of the pathogen population. However, each organ is associated with a specific and distinct probability of yielding clonal expansion, likely reflecting their underlying biological/compositional differences.

### Patterns of inter- and intra-organ dissemination

Comparisons of barcode frequencies between two organ samples can be used to quantify the relatedness between two pathogen populations in the host, effectively revealing inter- and intra-organ spread. This metric, known as genetic distance (GD), is small when two samples are highly related and large when samples are not related. Low GD values (~0.1-0.2) require that the same barcodes be relatively abundant (not simply present) in both samples. Spreading event(s) therefore must have occurred prior to sampling to allow for sufficient time to result in the same barcode(s) being highly abundant in both samples or be substantial enough to where a large number of bacteria are transferred. GD is also influenced by outliers, and so a different metric, called RD (“resilient” genetic distance), quantifies the number of barcodes that contribute to the similarity between two populations in a manner that takes into account all detected barcodes (25) (Figure 1A). For example, if both RD and GD are low, only a few barcodes contribute to the genetic similarity between samples. In contrast, if RD is high and GD is low, the samples are similar because they share many barcodes, which often occurs when samples closely resemble the inoculum through recent temporal separation. Therefore, RD provides a metric to distinguish between samples that are similar because they share few (low RD) or many (high RD) barcodes. RD can be normalized to the number of barcodes in a sample, and this “fractional” RD (FRD) is a directional metric that quantifies the extent to which shared barcodes are represented by all barcodes in the sample. FRD_A-B_ is low when the shared barcodes in sample A and sample B represent a small fraction of the total barcodes in sample B, and large when the shared barcodes represent a large fraction of the total barcodes in sample B. FRD_B-A_ is the reverse, where the shared barcodes are normalized to the total barcodes in sample A (Figure 1A).

To depict dissemination patterns across the entire animal, we display GD and FRD values as heatmaps, which permit rapid visualization of the degree to which samples are similar (Figure S6-S11). At 4 hpi, all organs possess highly similar bacterial populations that share many barcodes (Figure S6), as they have only recently been separated from the inoculum. After this initial capture of bacteria, genetic distance between organs generally increases over time, suggesting that the populations are not substantially disseminating and mixing, and that populations within organs are continually being antagonized by host defense processes, consistent with overall decreases in CFU.

In mice with expanded clones in the liver 1 dpi (e.g., Figure 2D-H), the dominant barcodes were not detected in other organs or other lobes of the liver, indicating that these expanded clones were locally confined (Figure S7 asterisks). Similarly, in animals with liver abscesses apparent 4 or 5 dpi, the bacterial populations in the lobe with the abscess were largely distinct from populations in other organs (Figure 3A, Figure S11). Dissection and comparison of individual abscesses from the same animal revealed that these populations were distinct (Figure 4), suggesting abscesses do not originate from the same bacterial population and do not substantially exchange bacteria. Surprisingly, multiple clones, rather than one, comprise a single abscess. In contrast to the liver, bacterial expansion events in other organs were associated with dissemination. The most significant spreading events were observed when ExPEC reached the bile (Figure 3B, Figure S10 (Mouse 17, 20), S11 (Mouse 31)). Each of the three animals with CFU in bile harbored a single highly dominant clone (>99% of all reads) and these clones spread systemically to multiple organs; however, the precise organs to which the clone disseminated varied across mice, suggesting that spread of these clones are stochastic (Figure S10-S11). Furthermore, FRD_organ-bile_ was significantly greater than the FRD_bile-organ_, confirming that systemic organs possess relatively more abundant non-transferred populations than bile (Figure 3C). More broadly, FRD values across all organs with some relatedness (GD < 0.8) decreased significantly over time, indicating that relatedness at later time points were the result of relatively few barcodes (Figure 3D, few barcodes relative to the total barcodes in a sample, not the entire library). Taken together, these results reveal that early genetic relatedness is driven by recent temporal separation (i.e., many barcodes), while later genetic relatedness is typically driven by the dissemination of a few clones.

**Figure 3.**
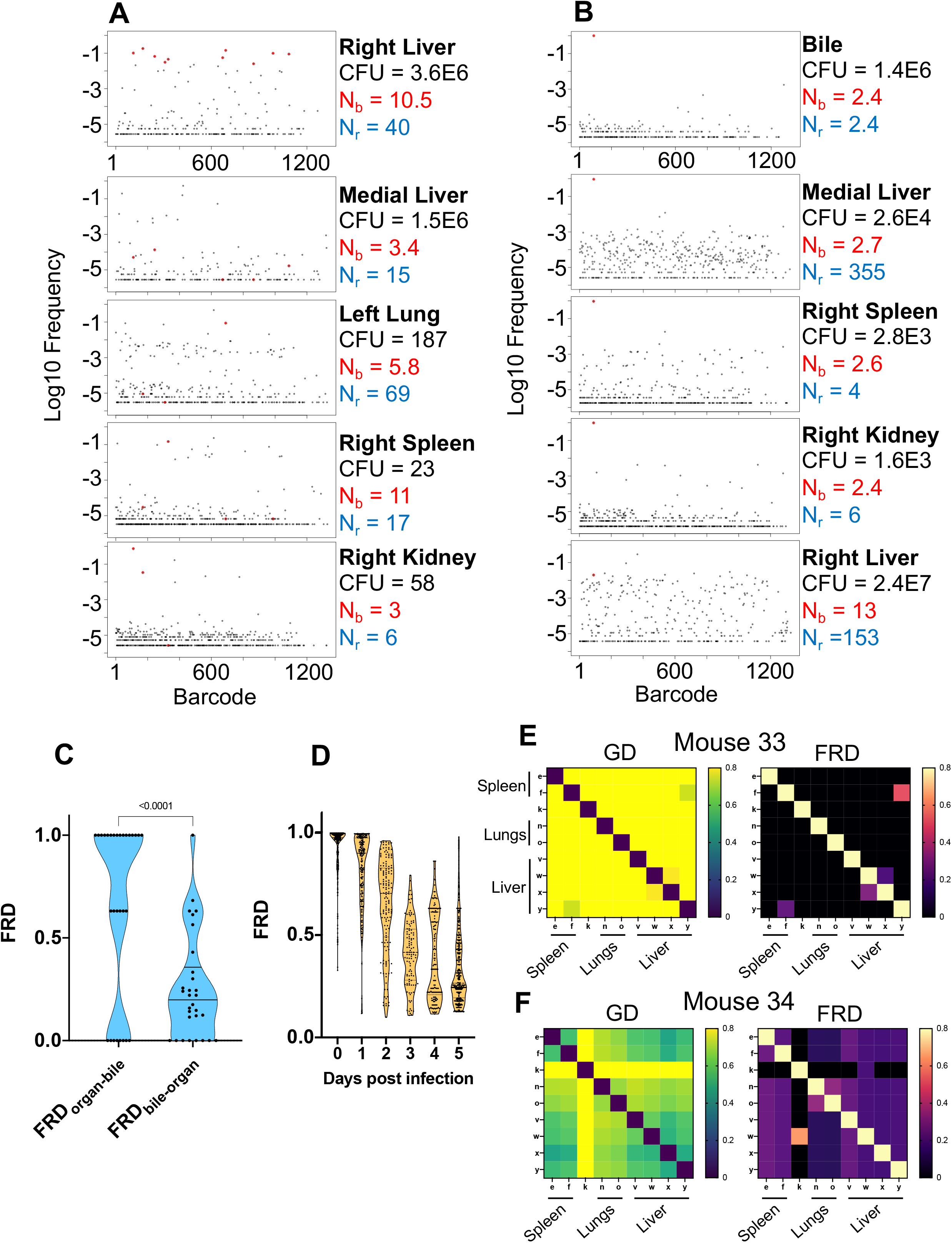
Visualizing dissemination patterns. A) Barcode frequency distribution from the right lobe of the liver in a single animal (mouse 24) is shown. This lobe harbored an abscess, which is reflected in the large bacterial burden. The top 10 barcodes are highlighted in red and identified in the four organs below. The relative absence of these dominant barcodes in the other organs reveals that the clones in the abscess did not substantially disseminate to other organs. GD and FRD values resulting from these comparisons are shown in Figure S11. N_r_ and N_b_ values are shown adjacent to each sample. B) Same as A but where the top barcode frequency distribution comes from the bile of mouse 20. The most dominant clone is highlighted in red and identified in other organs below. This clone is also dominant in other organs with exception of the right lobe of the liver, where the clones within an abscess are dominant. C) FRD values resulting from organ comparisons with the bile in all animals with bile CFU (Mice 17, 20, and 31). Significantly lower FRD values (Mann-Whitney) when the non-bile organ is set to the reference (FRD_bile-organ_) reveals that the non-bile organ harbors a more substantial non-transferred population, particularly exemplified by liver samples in panel B. D) FRD values for all organs, omitting FRD values of 0 (no detectable relatedness) and 1 (self-organ comparisons). This panel summarizes and quantifies the increasingly purple colors observed across Figure S6-S11. The significant decrease in FRD (Spearmans r = −0.7, p < 0.0001) indicates that relatedness becomes driven by fewer clones over time. E-F) Heatmaps of GD (left) and FRD (right) are displayed for mouse 33 (C) and mouse 34 (D). Lower GD values in mouse 34 indicates substantially more sharing of bacteria than in mouse 33. Low FRD values across all organs in mouse 34 reveals that only a few barcodes (i.e. clones) are being shared. FRD = 0 for most organs in mouse 33 because no clones are shared. In E-F, organs displayed are the right spleen (e), left spleen (f), left kidney (k), right lung (n), left lung (o), caudate liver (v), right liver (w), left liver (x), medial liver (y). Column names in FRD heatmaps represent the organ used as the reference. Heatmaps for all mice and all time points are shown in Figures S6-11.

**Figure 4.**
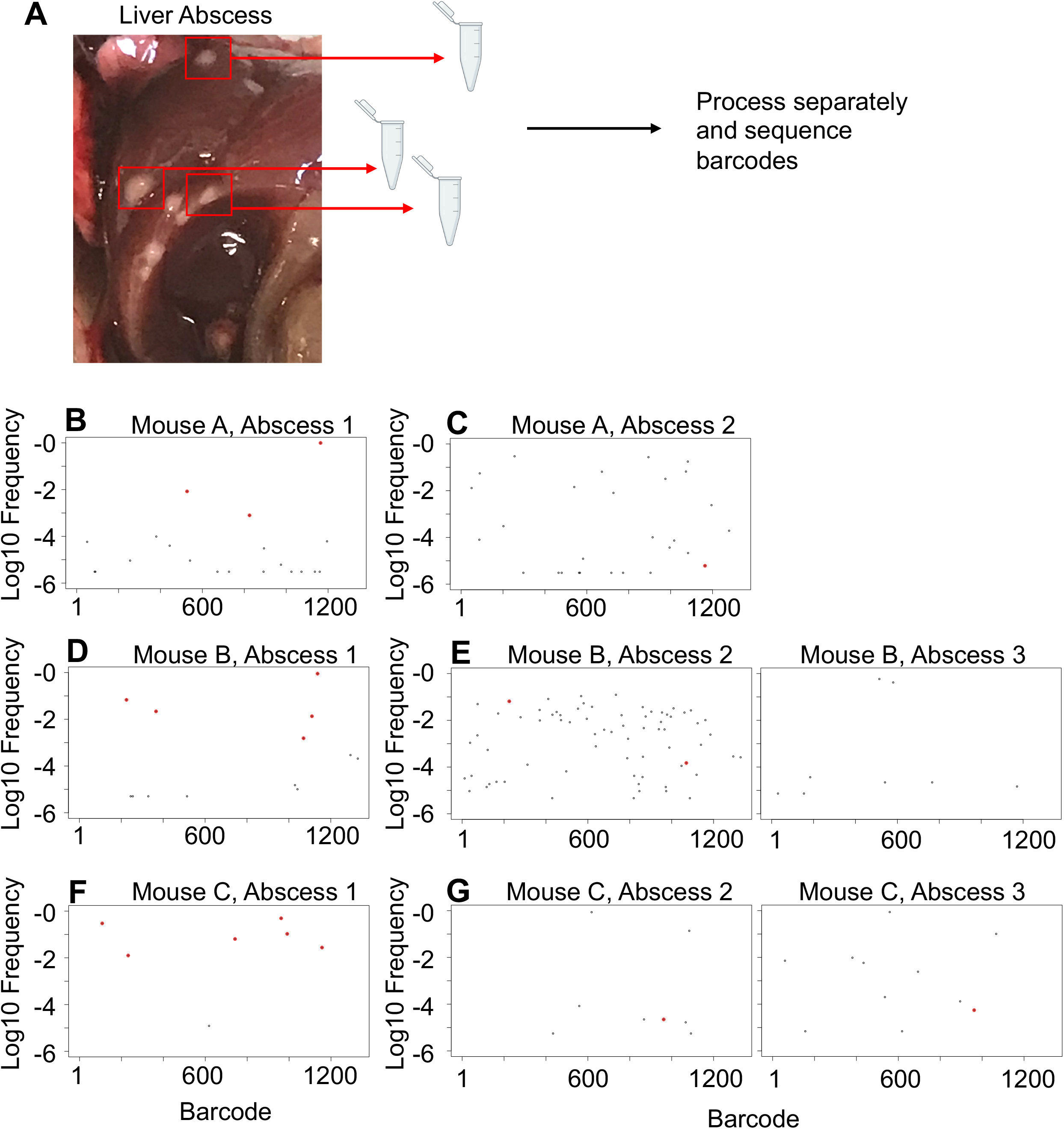
Liver abscesses arise from distinct bacterial populations. A) A schematic of the experimental setup is shown. Single abscesses were dissected and separately processed for analysis of barcode frequencies to determine if abscesses in the same animal arose from similar or distinct populations. The panels on the left (B, D, and F), where dominant barcodes are highlighted in red, correspond to the reference for the graphs on the right (C,E,G). These barcodes are then identified on the panels on the right (C, E, and G). Very few dominant red barcodes in C, E, and G indicate that populations within individual abscesses are distinct.

At higher resolutions, the heuristic value of heatmaps to visualize patterns of dissemination can be illustrated by comparisons of individual mice, such as mouse #33 and #34, littermate animals sacrificed 5 dpi (Figure 3EF). The GD-based heatmap identifies instances of transfer (bluer colors), and FRD-based heatmaps reveals the fraction of barcodes that contribute to transfer, a metric (in addition to FP) to gauge clonality. Mouse #33, despite having two liver lobes with abscesses containing 10^6^ bacteria, lacked any detectable spread between organs, indicated by the entirely yellow GD heatmap (Fig 3E, GD > 0.8). In contrast, mouse 34 had substantial systemic sharing (relatively blue GD heatmap, Fig 3F), where a single clone was dominant across all organs (blue/purple FRD heatmap, black colors indicate that no barcodes are shared). Bacterial burdens in the spleen and lungs were also 2-5 fold higher in mouse 34 than mouse 33. The presence of this clone highlights the substantial impact a single bacterium from the inoculum can have on pathogen distribution in the host. Thus, our approach enables detection and quantification of stochastic clonal expansion events; observing such events, which can remain localized or spread, deepens understanding of the origins of within host pathogen burdens. Collectively, our data reveal that clearance failures and systemic dissemination are mostly attributable to ~0.0001% of the inoculum. The manifestation of these events is highly variable across mice but can fall into three trajectories: 1) complete clearance without dissemination, 2) clonal expansion without dissemination (e.g., liver abscesses), and 3) clonal expansion with dissemination (e.g., bile)

### Host bottlenecks are widened by increased pathogen dose and macrophage depletion

The in-depth characterization of ExPEC systemic infection presented above served as a framework that enabled us to begin to investigate the factors and mechanisms that govern these dynamics. Many perturbations in infection models are known to alter bacterial burdens, although quantifying the degree to which changes in CFU result from wider (more permissive) bottlenecks and/or from increased replication has been challenging. The metrics employed in this study can be used to disentangle these processes.

We first assessed how bacterial burden and bottlenecks are influenced by infectious dose, which is fundamentally linked to infection outcome (33, 34). The barcoded ExPEC library was i.v. inoculated at two doses (5E6 and 2E7) and the livers and spleens were harvested at 5 dpi. Note that the relatively small difference in the sizes of these inocula also allowed us to assess the sensitivity of CFU and FP to modest differences in dose. Splenic CFU was highly consistent within groups, as observed above, and a 4-fold increase in dose resulted in ExPEC burdens ~2.5-fold greater than in the low dose group (Figure 5A). N_b_ and N_r_ in the spleen were also relatively similar within the groups, indicating relatively uniform barcode distributions, and both values were significantly larger in the high dose group, suggesting that there is a general correlation between dose, CFU burdens, and FP sizes in the spleen. Hepatic bacterial burdens were much more variable, consistent with data in Figure S4. Unexpectedly, there was a reduction in the bacterial burden in the liver at the high dose; a greater incidence of abscesses in the low dose group (6/10 animals) compared to the high dose group (2/9 animals) lead to a reduction in median CFU at the high dose. Despite the higher hepatic burdens in animals given the low dose, N_r_ was significantly higher in animals given the high dose, reflecting the role of clonal expansion in elevated CFU. The scaling of FP with dose reveals that a fixed fraction, not number, of bacterial cells from the inoculum survive clearance mechanisms up to 5 dpi. However, in contrast to the spleen, the dramatic replication of very few clones in the liver obscures this relationship from being detected by CFU alone.

**Figure 5.**
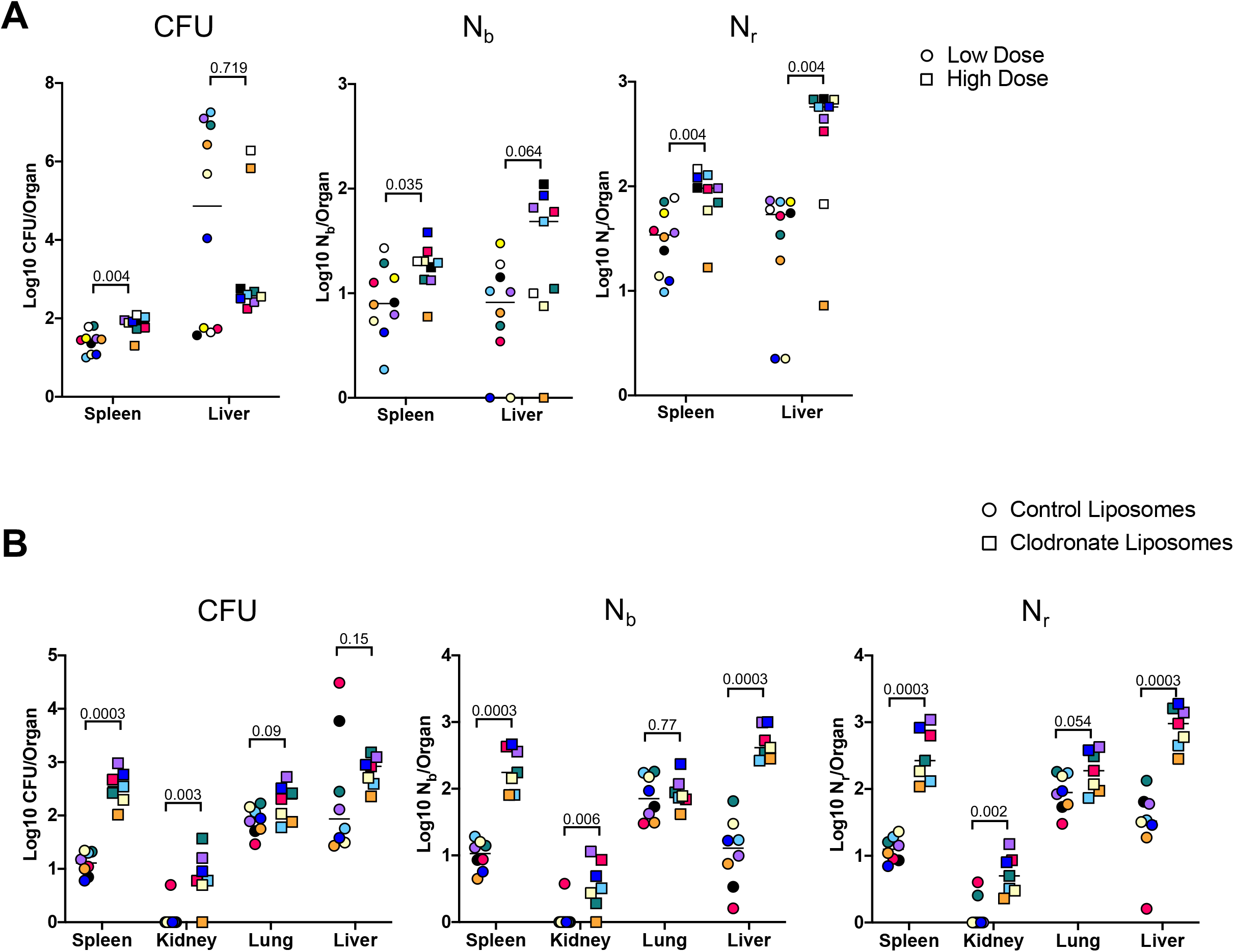
Inoculum size and macrophages regulate bottleneck sizes. A) CFU, N_b_, and N_r_ values from mice given a low (~5E6, circles) and high (~2E7, squares) dose of ExPEC i.v. 5 dpi. Note that all measurements in the spleen were significantly higher in mice inoculated with a higher dose, but only N_r_, not CFU, was significantly greater in the liver. Low FP values with high CFU indicate that a few clones replicated substantially. Points with the same color and shape are derived from the same animal. B) CFU, N_b_, and N_r_ values from mice treated with control liposomes (circles) or clodronate-containing liposomes (squares). CFU is significantly increased in the spleen and kidneys, but not the lung and liver. FP is significantly greater after clodronate treatment in the liver. Points with the same shape and color are from the same animal. The two animals with high CFU in the liver result from the expansion of very few clones (high CFU, low FP). Statistical significance for each comparison represents q-values from multiple Mann-Whitney tests.

To begin to assess the host mechanisms that control bottlenecks, we tested how bacterial burden and bottlenecks are influenced by tissue-resident macrophages, which rapidly sequester bacteria that enter the bloodstream.(2, 3, 35). To assess the contribution of these cells in establishing the bottleneck to systemic ExPEC infection, mice were treated with clodronate-containing liposomes, which depletes macrophages from the vasculature, particularly in the liver and spleen (36). The next day, mice were i.v. inoculated with ExPEC at the low (~5E6) dose used above. We chose a dose of clodronate and control liposomes that only partially depletes macrophages (37) because at higher, commonly used clodronate doses, animals rapidly succumb to infection (17). There was an increase in splenic CFU 5 dpi in animals treated with clodronate, whereas this difference was more subtle in the liver, lungs, and kidneys (Figure 5B). N_b_ and N_r_ calculations revealed that macrophage depletion increased founding population sizes in the liver, spleen, and kidneys, although the difference in the lung was not statistically significant. The significant increase in FP, but not CFU, after clodronate treatment in the liver is explained by two animals in the control group with the dramatic expansion of very few clones (Figure 5B, red and black circles). Since clodronate treatment increased both FP and CFU, macrophage depletion widens the bottlenecks in the liver, spleen, and kidney. Taken together with the dosing experiment, these observations underscore how stochastic expansion of clones can obscure detection of biologically significant signals, highlighting the utility of using barcoded libraries during experimental infection for even subtle changes in experimental parameters. In addition to being able to detect these signals, barcoding approaches can reveal whether elevation in CFU burdens are attributed to heightened bacterial replication within organs or wider host bottlenecks. In this model, increased dose and macrophage depletion widened bottlenecks in the host but did not increase the levels of bacterial replication.

### Bacterial genetic analyses disentangle organ-specific and temporal infection outcomes

Our STAMPR-based analyses uncovered several patterns of pathogen population trajectories in infected hosts, including eradication early during infection, survival/persistence until late in infection, and clonal expansion with or without dissemination. Moreover, each of these trajectories had organ-specific patterns. However, since our analyses relied on isogenic bacterial strains, we were unable to infer the pathogen factors that contribute to these distinct outcomes. We used transposon-based genetics to begin to uncover these factors. We reasoned that identification of bacterial genes required for early survival, but not for subsequent expansion or persistence, would 1) signify that early and late survival/expansion are at least in part separable and 2) identify the pathways that distinguish them. To identify early pathogen survival determinants, we analyzed an ExPEC transposon library composed of ~150,000 unique transposon mutants 1 day after i.v. inoculation. As expected (Figure S4, 2EG), we detected highly dominant mutants (i.e., clones) in the livers and spleens of 3/5 mice (Figure S12). Since these three animals also had markedly high bile CFU, these clones are likely also present in the bile.

Comparisons of the abundances of gene insertion frequencies in samples isolated from the liver and an input that was replated on LB were carried out to identify processes that contribute to early ExPEC survival using a new analytic pipeline (see methods) that simulates infection bottlenecks. There was a predominance of cell envelope homeostasis genes and gene clusters that were underrepresented in liver samples, including *mlaFEDCB* (38, 39)*, ftsEX* (40, 41)*, envZ/ompR* (42), and many LPS biosynthetic genes (Figure 6A), suggesting that cell envelope homeostasis across multiple pathways is required for extraintestinal survival. Another coherent set of underrepresented genes in the liver included *manA, nagAB, pgi*, which convert mannose-6-phosphate, n-acetylglucosamine-6-phosphate (GlcNAc-6P), and glucose-6-phosphate into fructose-6-phophate respectively (43) (Figure 6B). GlcNAC-6P, the substrate of NagA, is generated in *E. coli* through two mechanisms (44). In the first, extracellular GlcNAc is phosphorylated and imported through the mannose phosphotransferase system. In the second, GlcNAc is released as a peptidoglycan degradation product, and is phosphorylated by NagK (GlcNAc kinase). Interestingly, *nagK* was not underrepresented in the liver, suggesting that the host is the source of GlcNAc. Taken together, the screen revealed that several cell envelope homeostasis pathways are important for early survival in the liver and an unexpected requirement for fructose-6-phosphate during systemic infection.

**Figure 6.**
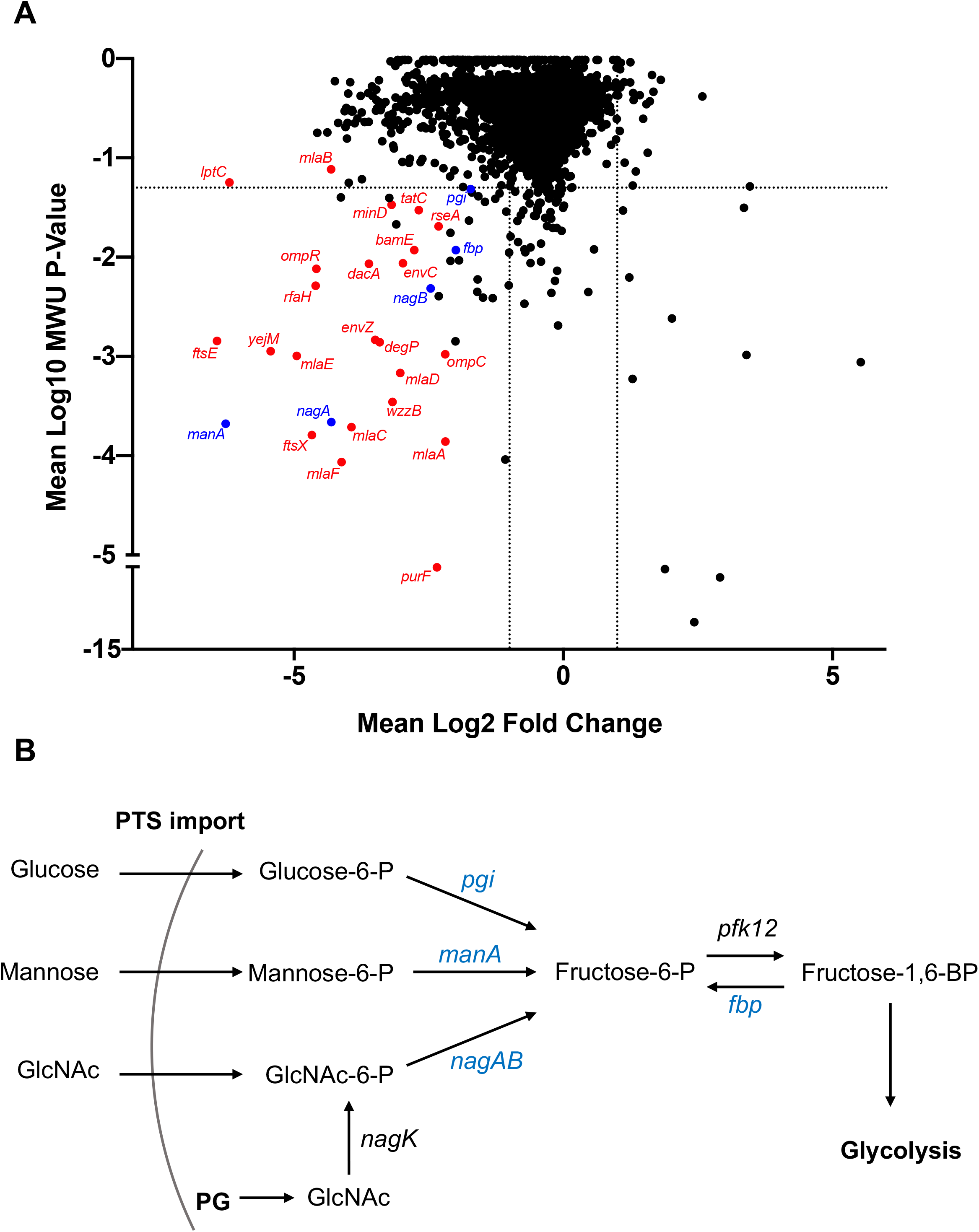
Genes that promote ExPEC survival in the liver. A) Inverse volcano plot of fitness of ExPEC genes in the liver. Genes shown in red are involved in cell envelope homeostasis and genes shown in blue are involved in generating fructose-6-phosphate, and their roles are schematized in B. B) Pathways to convert sugars to fructose-6-phosphate. GlcNAc generated from peptidoglycan (PG) degradation is converted to GlcNaC-6P by *nagK*, which is not among the underrepresented genes in A.

To investigate how the genetic requirements for ExPEC survival vary during infection, we chose a subset of the most underrepresented genes/pathways for early hepatic survival to test for survival across organs and over time. Traditionally, competition between WT and mutant strains are performed. These studies rely on CFU plating to determine the relative abundance of the mutant strain before and after infection (45–48). However, as we show above, this approach would likely be confounded by early stochastic clonal expansion events (particularly in the liver). Instead, we adopted a barcoding strategy similar to previous studies (47, 49). In-frame deletions of *mlaC* (RS18735)*, ftsX* (RS20130)*, yejM* (RS12925)*, manA* (RS09455)*, nagA* (RS03530), and *nagK* (RS06530) were created and each strain, along with the WT, was barcoded with 3 tags and pooled, resulting in an inoculum comprised of a mixture of 21 tags. The Δ*nagK* strain served as a “negative control”, as it was not underrepresented in our screen (Table S1). This library was i.v. inoculated and the bile, spleen, right kidney, lungs, and all 4 lobes of the liver (separately) were harvested 1 and 5 dpi. Barcodes in each sample were then sequenced to determine the abundance of all 21 tags, and therefore each mutant.

One dpi in the liver, stochastic clonal expansion was observed and manifest as a dramatic increase in the abundance of a few barcodes and the corresponding marked decrease in the abundance of all other tags in the sample, even those from the same genotype. Thus, clonal expansion explains the large variance in fitness measurements. For example, one animal had a substantial expansion of one of the three tags barcoding the *mlaC, manA*, and *nagA* mutants in the caudate lobe of the liver (Figure 7A, red asterisks), within an associated drop in the abundance of all other barcodes from the animal, even the other barcodes marking the same mutants. In the liver, all genes found to be underrepresented in the transposon screen displayed fitness defects. However, the magnitude of the defects in the *mlaC, ftsX, yejM*, and *nagA* mutants were far more pronounced than that observed in the *manA* mutant, which possessed modest but consistent defects relative to the WT. Clonal expansion events likely prevented *manA* from reaching statistical significance (Figure 7A). Similar patterns of mutant fitness defects were observed in the spleen and the kidneys, although *manA* was not statistically different than the WT 1dpi in the kidneys (Figure S13). Intriguingly, there was no difference between Δ*manA* and Δ*nagA* compared to Δ*nagK* and WT in the lungs, indicating that these genes have organ specific importance for ExPEC survival and reveals that the lung microenvironment has particularly distinct selective effects (Figure 7C). The Δ*nagK* negative control was identical to the WT across all organs 1 dpi. The differential importance of *nagA* and *nagK* confirms our transposon results and provides further evidence that the source of GlcNAc-6P is the host.

**Figure 7.**
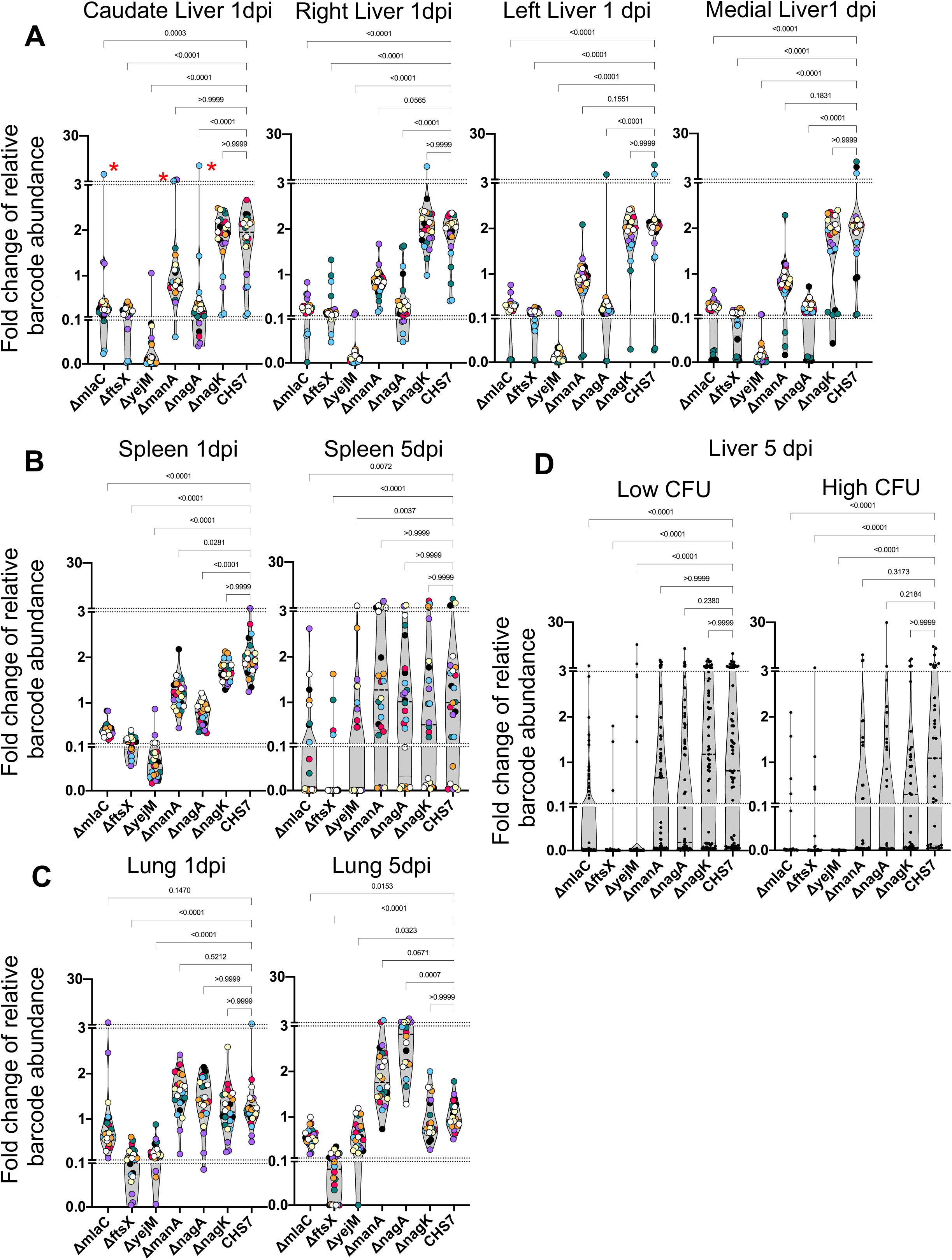
Distinct genetic requirements for ExPEC survival/growth across organs and time. A) Changes in barcode frequencies for WT and 6 mutants after coinfection are shown for liver 1dpi. Red asterisks represent an example of clonal expansion. B and C) same as A) for spleen (B) and lungs (C) 1 and 5dpi. D) Same as A-C but for liver samples 5 dpi, separated into lobes with or without abscesses. Figure S13 displays this data separated by lobe, in addition to data from the kidneys. Within each graph, points with the same color were derived from the same animal. Each strain is represented by 3 barcodes, so within each graph, each color is represented 21 times. Dotted lines depict breaks in the y-axis that are useful to visualize the influence of clonal expansion. P-values (Dunn’s multiple comparisons test) are shown relative to the WT.

We then assessed the role of these genes for survival at 5 dpi, during the persistence and expansion phase. In this cohort, 5/8 animals had at least one lobe of the liver with an abscess, and therefore several orders of magnitude more bacteria than lobes without abscesses (Figure S13). Since each liver lobe was segregated, however, we grouped lobes with and without abscesses separately for analysis; this grouping enables assessment of genetic requirements for persistence (no abscess) or for expansion (abscess). In addition, we omitted the kidneys from this analysis, as too few bacteria were recovered 5 dpi (Figure S13).

The Δ*mlaC*, Δ*ftsX*, and Δ*yejM* strains were significantly depleted in the spleen, liver (with or without abscesses) and lungs relative to the WT at 5 dpi, further highlighting the essentiality of the cell envelope-related process to survival within these organs (Figure 7B-D). In contrast, despite having clear defects at 1 dpi, Δ*nagA* and Δ*manA* were indistinguishable from WT and Δ*nagK* in the spleen and the liver at 5 dpi, regardless of the presence of abscesses. An intriguing comparison is Δ*nagA* and Δ*mlaC*, which display nearly identical fitness defects in the liver 1 dpi but clearly differ by 5 dpi (with or without abscesses). To our surprise, Δ*nagA* was significantly more abundant in the lungs than the WT or Δ*nagK* at 5 dpi (Figure 7C). Therefore, cell envelope and sugar metabolism genes likely contribute to distinct mechanisms that facilitate ExPEC survival, in the face of host factors that fluctuate over time and between organs. These results further reveal that the pathogen factors that are required for the initial establishment of infection differ from those required to survive later or clonally expand. Specifically, ExPEC requires hexose metabolism to establish populations within specific organs, but this pathway is not required by the sub-populations that ultimately persist later in the infection or clonally expand, suggesting that establishment, persistence and expansion depend on distinct pathogen pathways. These data further highlight how understanding within host infection dynamics provides critical contextualization of pathogen virulence or fitness factors that vary in their importance across time and space.

## Discussion

Here, using pathogen barcoding and a high-resolution approach for comparing barcode frequency distributions, we describe the complex infection dynamics of systemic ExPEC infection (summarized in Figure S14). Although we found that ExPEC is largely cleared by most organs, we observed organ-specific population dynamics and discovered that clearance failures are pervasive, particularly in the liver. These failures manifest as dramatic expansions of very few bacterial cells from the inoculum. Some expansions are not easily revealed by CFU-based metrics (e.g. clonal expansion at 1 dpi in the liver), but our analytic approach readily detects and quantifies these events in addition to contextualizing them with populations at other organs. During experimental perturbations, our analyses also readily distinguished when increases in bacterial burden are due to widening of bottlenecks or to increased bacterial replication. We found that, despite clonal expansion, host bottlenecks were highly consistent between animals across all organs. However, bottlenecks are highly sensitive to subtle organ-specific changes in experimental parameters, including inoculum size and macrophage depletion. Furthermore, we identify a potential role for bacterial sugar metabolism that distinguishes early extraintestinal survival with later persistence or clonal expansion. Our analysis of population dynamics can also refine transposon-based studies of genetic requirements by contextualizing bacterial genes required for infection in the setting of temporal, organ-specific, and stochastic infection outcomes. Furthermore, understanding the population dynamics of an infection model facilitates the optimization of experimental designs in infection studies.

Our findings highlight the differential capacities of host organs to clear bacteria. In general, most organs continuously restricted the pathogen population over time, indicated by decreasing N_r_ values over time. A notable exception to this trend was the lungs, where FP values largely plateaued by 2 dpi, explaining why this organ had among the highest FP sizes 5 dpi. In addition, clodronate had no effect on CFU in the lung and some mutants defective for survival in the liver were more abundant than WT in the lung. These three lines of evidence indicate that the lung microenvironment is less hostile to ExPEC, perhaps reflecting distinct cellular niches occupied by ExPEC in the lungs relative to the liver or spleen. Furthermore, we discovered that ExPEC forms visible abscesses in liver at much higher frequencies than expansion events in other organs. Thus, the liver and lungs appear to differ markedly in their clearance defects; the liver appears more permissive for ExPEC replication, whereas the lung appears more permissive for bacterial survival.

Very few studies to date have reported on the population dynamics of systemic infections. In one study, i.v. inoculation of a *S. aureus* pool consisting of three differentially marked strains revealed several trends observed here (17). Although the use of 3 markers limited the resolution, this study of *S. aureus* dissemination also found that bottlenecks (measured as the proportion of organs with less than 2 strains) widened in size with infectious dose and macrophage depletion. However, unlike ExPEC where expansion of clones in liver abscesses remained locally confined, *S. aureus* dissemination from the liver was thought to result in clonal abscesses in the kidney. In general, abscess formation is much more consistently observed for *S. aureus* (50, 51) whereas this outcome is apparently stochastic in this ExPEC model. Thus, pathogen-specific factors exert distinct roles in the determination of the organ-specific outcomes of clonal expansion events. Another study examining systemic *P. aeruginosa* population dynamics with STAMP found that the pathogen consistently replicated in the bile and these cells became the source of bacteria in the intestinal tract (16). In ExPEC, the most dramatic instances of dissemination also occurred in animals with ExPEC in bile. The FP of the bile population in our study was ~1, whereas we calculated that the *P. aeruginosa* FP was ~10 times greater (25). The more restrictive bottleneck for ExPEC potentially explains why transit to the bile is stochastic with this pathogen. The mechanisms by which pathogens such as ExPEC and *P. aeruginosa* spread systemically from bile require further investigation but may be linked to the event(s) by which the pathogen seeds the bile to begin with.

The bacterial genetic determinants of systemic infection are of great interest, as these may inform development of therapeutics for the treatment of these severe infections (5, 48, 52–54). In models of overwhelming bacteremia, these genetic determinants often encode surface structures, such as the capsule in K1 ExPEC (55). Similarly, we found that most of the underrepresented genes in the liver and spleen encoded components of the cell envelope. Unexpectedly, there was little overlap between genes we identified to be underrepresented in the liver or spleen and genes identified as underrepresented in a previous study using a similar experimental system (48). A potential explanation for this discrepancy is that Subashchandrabose et al performed their experiments in a different mouse background (CBA/J), and different strains of mice impose distinct barriers to infection, altering the pathogen genetic requirements for infection (56). In addition, it is unknown whether clonal expansion occurs in CBA/J mice. Our findings suggest that understanding population dynamics prior to determining genetic requirements can refine identification of genetic dependencies in the presence of genotype-independent expansion events.

Analysis of barcode frequency distributions permits a more quantitative and deeper dissection of population dynamics in experimental infection models beyond enumerating bacterial burdens within organs. The 8 metrics we used (Figure 1A, (25)) to compare pairs of organs within an animal allows for delineation of patterns of expansion and dissemination. For example, we show how some expanded clones (i.e., in liver abscesses) appear to have relatively little influence on barcode frequencies in other organs in the animal (Figure 3A). In stark contrast, Figure 3B reveals an example in which a single clone in the bile profoundly alters the pathogen distribution in multiple organs. In many cases, we observed that CFU values were similar to FP values (Figure 1I, Figure S2-S4), suggesting that the respective bacterial population had not undergone substantial net replication. Heatmaps enable direct visualization of dissemination patterns and the degree to which these dissemination events manifest within an organ. Heatmaps from animals 5 dpi revealed marked heterogeneity in dissemination patterns that vary across organs (Figure S11). Furthermore, these metrics reveal the variability between animals both in terms of clonal expansion within organs, bottlenecks between organs (Figure S2-S4), and dissemination within and between organs (Figure S11). Beyond comparisons between animals, our metrics can be applied to single animals and revealed that stochastic expansion and dissemination of very few cells from the inoculum was pervasive.

Early stochastic clonal expansion events were detected in every experiment in our study, using three independent libraries and distinct approaches (STAMPR, transposon libraries, and barcoded mutants). These expansion events underly variation in infection outcomes. While dissection of the mechanism(s) that account for this stochasticity is beyond the scope of this study, our observations suggest several factors account for and modulate clonal expansion. Such events appear to fall into two functional categories – those that are due to dissemination (e.g. from bile) and those that are not (e.g. liver abscesses). In the latter, we observed an apparent inverse correlation with infectious dose, although more animals are required to reach statistical significance (Figure 5A). This trend suggests that more robust early immune induction (presumably elicited with a higher dose) disproportionately clears bacteria, a hypothesis that has also been generated by mathematical modeling (57, 58). Additionally, we observed apparent lobe (more frequent in the right lobe) and topological (all on the surface) biases in liver abscess formation, raising the possibility that anatomic differences between lobes and their relative locations contribute to apparently stochastic outcomes, as some lobes may be more likely to be seeded (due to blood flow) and more permissive to bacterial replication. Furthermore, clonal expansion explains why bacterial burden is highly variable in EXPEC and may also explain high variability in systemic infection outcomes in other bacteria (59).

The STAMPR analytic framework (25) applied here to study infection population dynamics revealed how very few cells in a large inoculum disproportionally contribute to pathogen burdens in different host organs. By monitoring patterns of dissemination, we uncovered that these few cells can drastically alter pathogen distributions in distal organs. Our findings suggest that perturbations that impede the expansion of the few cells that clonally expand could profoundly alter the course of infection. Finally, our observations of organ-specific pathogen population dynamics and genetic requirements suggests that investigation of host tissue-specific factors that regulate these dynamics, perhaps centered around bacterial metabolism of host-derived sugars, may uncover novel facets of host-pathogen interactions.

## Methods

### Ethics statement

All animal experiments were conducted in accordance with the recommendations in the Guide for the Care and Use of Laboratory Animals of the National Institutes of Health and the Animal Welfare Act of the United States Department of Agriculture using protocols reviewed and approved by Brigham and Women’s Hospital Committee on Animals (Institutional Animal Care and Use Committee protocol number 2016N000416 and Animal Welfare Assurance of Compliance number A4752-01)

### Media and antibiotics

Bacteria were routinely cultured in LB at 37°. Additional compounds and antibiotics were used at the following concentrations: kanamycin 50 μg/ml, chloramphenicol 20μg/ml, carbenicillin 50μg/ml, diaminopimelic acid (DAP) 300μM. Bacteria scraped from plates were resuspended in PBSG (PBS 25% glycerol)

### Animal experiments

All animals used in this study were 8–10 week old female C57BL/6J mice obtained from Jackson Laboratories and housed for 3 days prior to handling. Defined volumes of frozen cultures were thawed and resuspended in PBS for i.v. inoculations. Prior to inoculation mice were gently warmed on a heating pad and restrained in a Broome style restrainer (Plas Labs), and 100 μl of bacteria was injected into the lateral tail vein with a 27G needle. Cages were then left undisturbed until the time of sacrifice. For clodronate (or control liposome) treatment, 50μl was injected i.v. 1 day prior to infection. For all experiments in the study, the bile in the gallbladder was aspirated with a 30G needle into 100 μl of PBS, after which the gallbladder was surgically removed prior to harvesting the liver. We detected bile CFU in 4 animals across all experiments in this study. Three are detailed in the first experiment (Figure S2, S10-S11) and the fourth was present in the clodronate-treated cohort but was moribund prior to the time of sacrifice and was therefore excluded from analysis. The “left lung” samples described in the first experiment (Figures S2-S4) includes the post-caval lobe. All “blood” samples are derived from a 200 μl cardiac bleed. After sacrifice, organs were homogenized with two 3.2mm stainless steel beads (Biospec) for 2 minutes in PBS and plated on LB + kanamycin plates.

### Bacterial strain construction

CFT073 *rpoS*^-^ (29, 31, 60) and CFT073 *rpoS^-^* Δ*mlaC* were conjugated (500 μl donor with 500 μl recipient overnight on a 0.45 μM filter) with *E. coli* donor strain MFDλpir (61) harboring a pDS132 (62) derivative containing appropriate homologous recombination fragments to facilitate repair of the *rpoS* allele. Transconjugants were selected on LB + chloramphenicol, re-streaked on LB + chloramphenicol, and cultured overnight with chloramphenicol. 300μl was diluted in 3 ml of LB with 10% sucrose, cultured overnight at 37°, and plated for single colonies on LB with 10% sucrose. Sanger sequencing was used to identify colonies with the repaired *rpoS* allele. Positive colonies were further confirmed by whole-genome sequencing through MiGS (Pittsburgh, Pennsylvania, USA). The CFT073 strain with the corrected *rpoS* allele is designated as CHS7.

The same sucrose-based allele exchange method was used to create in-frame deletions of *manA, yejM* (C-terminal domain only), and *ftsX* from CHS7. For deletion of *nagA* and *nagK*, we adopted a CRISPR-based counterselection approach used in (63) with several modifications. We first introduced *bla* and *sacB* from pCVD442 (64) into pSU-araC-Cas9, creating plasmid pCAS, which contains an arabinose-inducible *cas9*. A separate plasmid containing the guide RNA was generated from pTargetF (63) by introducing homology fragments in addition to *sacB, cat*, and conjugation machinery from pDS132, creating a family of plasmids referred to as pGuide. Derivatives of pGuide were created with the appropriate spacer targeting the gene of interest and homology fragments. To generate deletions in a target gene, the recipient strain was first transformed with pCAS, which harbors a beta-lactamase gene conferring carbenicillin resistance. pGuide was then introduced into a pCAS-carrying recipient strain by conjugation and transconjugants were selected on plates containing chloramphenicol, carbenicillin and 0.2% arabinose overnight at 37°C. The transconjugants were then scraped and re-plated on plates containing chloramphenicol, carbenicillin and 0.2% arabinose overnight at 37°C. Single colonies from this second passage were streaked again on plates containing chloramphenicol, carbenicillin and 0.2% arabinose. Single colonies from this passage were then streaked on LB plates containing 10% sucrose and lacking NaCl to counter-select both plasmids, which was confirmed by loss of the resistance markers. This protocol was initially optimized by targeting *lacZ*, where blue/white screening could be used to determine the steps at which successful edits have been made. Successful deletions were confirmed by PCR and, after a subsequent colony purification, a second PCR. In addition, both the *nagA* and *manA* mutants were unable to grow in minimal media supplemented with GlcNAc or mannose as the sole carbon source, respectively.

### Construction of barcoded ExPEC library

The plasmid used to introduce barcodes into CHS7 was created by amplification of a kanamycin fragment adjacent to *lacZ* with a primer containing 20 random nucleotides (Integrated DNA technologies). This fragment was cloned into pDS132 using the NEB HiFI DNA Master Mix (New England Biolabs) and transformed into MFDλpir and plated on LB+ kanamycin + DAP, yielding pDS132-STAMPR. The cloning reaction was scaled up to yield 70,000 colonies. Ten colonies were individually Sanger sequenced to confirm the presence of a single, unique barcode. Cells were then frozen into individual aliquots.

To introduce barcodes into CHS7, 30μl aliquots of the barcode donor were thawed in 3 ml LB + kanamycin + DAP. A 1:1 ratio of the donor and recipient CHS7 were mixed, pelleted, and resuspended in 100μl of PBS. Suspensions were spotted on 0.45 μm filters and incubated at 37°C for 3 hours, after which the cells were resuspended in PBS and plated on LB + kanamycin. Barcodes in 10 individual colonies were Sanger sequenced to confirm that each recipient had a single distinct barcode. Finally, 1152 individual colonies were inoculated in twelve 96-well plates in LB + kanamycin, grown overnight, pooled, and frozen in PBSG

### Barcode stability and influence on ExPEC growth

To assess if the integrated pDS132-STAMPR is maintained in the absence of selection, we cultured the library in the absence of kanamycin overnight. 1/1000 dilutions of the culture were serially passaged overnight for 5 days and plated daily on LB with or without kanamycin. Additionally, 30 colonies each day were patched from the plate lacking kanamycin onto LB + kanamycin. All patched colonies on all 5 days were resistant to kanamycin, and thus possessed the barcode. To assess if the pDS132-STAMPR confers an *in vitro* growth defect, 9 different barcoded strains along with CHS7 were growth overnight diluted 1/1000 and individually grown in triplicate in a 96 well plate overnight. OD600 was measured in a plate reader.

### Sample processing for sequencing

Bacterial samples obtained from either *in vitro* culture or organ homogenates were plated on LB + kanamycin plates and incubated overnight at 37° C. If the cultures yielded a high density of cells, they were scraped and resuspended in PBSG and aliquots were stored at −80° C. If the cultures yielded few cells, each colony was individually picked and resuspended in PBSG. Suspensions were frozen at −80°C until processing. Frozen aliquots of cells were later thawed and 2 μl was diluted in 100 μl of water. Samples were then boiled at 95°C for 15 minutes and 1μl was used as template for PCR. Cycling conditions for PCR were the following: Initial denaturation 98°C 30s, 25 cycles of 98°C 15s, 65°C 30s, and 72°C 15s, and final extension of 72°C 5 min. Reactions were carried out with a total volume of 50ul using Phusion DNA polymerase (New England Biolabs). Amplified products were then run on 1% agarose gel to verify the presence of the amplicon and all samples were pooled and purified using the GeneJet PCR Purification Kit (Fisher). Purified amplicons were quantified by Qubit and loaded on a MiSeq (Illumina) with either V2 or V3 reagent kits for 78 cycles. FASTQ files were then processed as described in (25) to yield STAMPR measurements (N_b_, N_r_, GD, RD, FRD)

### Creation of the STAMPR calibration curve and barcode reference list

After the barcoded library was pooled, 100 μl of 10-fold serial dilutions were spread on LB + kanamycin plates, corresponding to 10^1^ cells to 10^8^ cells. These samples were used to create a standard curve and serve as controls of known bottleneck sizes. All samples were prepared and sequenced as described above. Two replicates of the most undiluted sample were sequenced and the dedupe plugin on Geneious (Biomatters) was used (k=20, max substitutions = 2, max edits = 2) to deduplicate the reads to create a preliminary list of reference barcodes. Then, the two replicates used to create this list were combined with a third and mapped to the preliminary list of references in CLC Genomics Workbench (Qiagen) using the default mapping parameters. We then determined a depth threshold that corresponded to 97% of all reads. This was used to determine the set of 1329 barcodes that was used throughout this study. Note that this is slightly greater than the number of colonies picked (1152) thus providing a “cushion” against errors in deduplication or mapping. This list of references was then used to map all samples of the calibration curve. The output vector containing the number of reads mapped to each barcode was then processed for data analysis as described in detail in (25), to confirm that this list of references yields accurate FP estimates (Figure S1C).

### Creation of ExPEC transposon library

500μl of E. coli donor MFDλpir containing the transposon bearing plasmid pSC189 (65) grown overnight was mixed with 500μl of CHS7 grown overnight. The mixture was pelleted, resuspended in 100μl PBS and spotted on 0.45 μm filters and incubated overnight at 37°C. This procedure was performed in triplicate and tittered on LB + kanamycin plates to quantify transconjugants. For the final library, ~7 million transconjugant colonies were obtained and pooled.

### In vivo ExPEC transposon insertion sequencing (TIS) protocol and reads processing

2.5μl of the CHS7 transposon library was resuspended in 1 ml PBS, corresponding to 2E8 CFU/ml. 100μl of this suspension was injected i.v. into 5 mice. The next day, the liver and spleens were harvested, homogenized, and plated on 245mm or 150mm LB + kanamycin plates, respectively. Cells were scraped in PBSG and stored in 1ml aliquots. gDNA was then harvested using the GeneJet gDNA Isolation Kit. Generation of libraries for sequencing was performed essentially as previously described (47). Briefly, 10 μg of gDNA was fragmented to 400bp fragments with an ultrasonicator. Ends were then repaired with the Quick Blunting Kit (New England Biolabs) and A tailing was performed with Taq polymerase. Adapter sequences were ligated overnight with T4 DNA ligase. 29 cycles of PCR were then performed to enrich for transposon-containing sequencing, and an additional 19 cycles of PCR were performed to introduce sample indexes and Illumina sequences. 300-500bp fragments were then gel excised, quantified, and loaded on a MiSeq for 78 cycles.

All steps to generate BAM mapping files were performed using CLC Genomics Workbench. First, FASTQ files were trimmed using the following parameters. 3’ sequence – ACCACGAC; 5’ sequence - CAACCTGT; mismatch cost = 1; gap cost = 1; minimum internal score = 7; minimum end score = 4; discard reads < 10 nt. Trimmed reads were mapped to the CFT073 genome using the following parameters. Mismatch score = 1; Mismatch cost = 1; Insertion cost = 3; Deletion cost = 3, Length fraction = 0.95, Similarity fraction = 0.95. Reads that mapped to multiple locations were mapped randomly once, and a global alignment was performed to ensure accurate positions of terminal nucleotides. The resulting mapping file was exported in BAM format and processed as described below.

### RTISAn pipeline for TIS data analysis

We applied our new bottleneck calculation methodology (25) to refine our previous TIS data analysis pipeline (Con-ARTIST) (66), as it did not sufficiently account for infection bottlenecks. This new R-based TIS analysis (RTISAn) employs the resampling strategy used here to calculate FP (25) to accurately simulate infection bottlenecks from input TIS libraries. RTISAn serves as a successor to Con-ARTIST (66–68). The RTISAn approach is functionally similar to the “resampling” method used in the TRANSIT pipeline but is implemented in R (69). The RTISAn pipeline begins with the user input of a FASTA file and locations of annotations or features. A “TAinfo” file is generated that identifies the overlapping feature of each TA site. The TAinfo file is then used with a BAM mapping file to count the number of reads at each TA site, compiling this information in a “TAtally” file.

The bulk of the RTISAn computation is used to compare two TAtally vectors, where one is an input and the other is an output. The goal is to identify genes for which insertions are either under- or over-represented in the output relative to the input. First, both the input and output vectors are normalized via a sliding window approach for replication bias. We set a window and step size of ~100,000 nt, so ~50 windows will be made for the *E. coli* genome. The mean of the bottom 99% of insertions is used to normalize each window, preventing the effects of clonal expansion from influencing the normalization.

Next, the normalized input vector must be resampled to match the saturation of the output vector. To accomplish this, the normalized input vector is iteratively resampled from a multivariate hypergeometric distribution 100 times, starting from 1000 reads and ending at the number of reads in the normalized input vector. The saturation is plotted against the sampling depth; the sampling depth that best approximates the saturation in the output vector is determined with inverse linear interpolation. The output represents a single sampling depth that accurately simulates the bottleneck on the input such that the saturation matches the output. To refine this estimate, we take an similar approach used to identify local minima in (25); instead of identifying local minima, we resample the input vector for 200 iterations at various sampling depths such that the resampling yields saturations closer to the saturation of the output vector than the previous sampling depth. In doing so, 200 sampling depths are acquired, from which a mean and standard deviation is obtained. These values are used to sample a normal distribution 100 times. These 100 values represent read depths that when used to resample the input vector result in a saturation that approximates the saturation of the output vector. The final 100 simulations are performed at these read depths on the input vector.

Each simulation is compared on a gene-by-gene basis to the normalized output vector. We calculate a geometric mean Mann Whitney P value and mean fold change across all simulations. We also determine the number of “informative sites”, defined as the number of TA sites which are nonzero across the output and all simulations. Genes with few informative sites in the input will usually not pass significance cutoffs but can still display large fold changes.

Since both P value and fold change are separate informative metrics, a script that combines both sets of information to “rank” genes when multiple animal replicates are used was created. This approach is partly unbiased, in that the user supplies a desired P-value and fold-change cutoff for significance but the relative position of each gene with respect to the entire dataset is considered as well. The rank is first determined by creating a score, from 0 to 1, that measures how many instances a gene passed either the P-value or fold change (with cutoffs of 0.05 and −1 in this study, respectively). For example, if 5 animals are used, then a score of 0.9 implies that, out of 10 metrics (5 fold changes and 5 p-values), 9 passed the user-specified cutoffs. A separate unbiased rank is created by the mean rank of the average P value and fold change for each gene. The final rank is then generated from the mean of the unbiased rank and the score. In practice, this strategy ensures that genes that consistently pass the user specified cutoff will always be ranked above genes that pass the cutoffs fewer times; this latter category can then subdivided into the relative magnitude of fold change and P-value.

The entire RTISAn pipeline are available at https://github.com/hullahalli/stampr_rtisan including TAtally files to reproduce the comparisons in this study.

### Analysis of barcoded mutant libraries *in vivo*

Barcoding of each mutant was performed by conjugation of the same donor library used to generate the CHS7 barcoded library. Three colonies per mutant were Sanger sequenced to establish the barcode identity, cultured overnight, and pooled to generate a mixed library of 21 tags in 7 strains (*mlaC, yejM, ftsX, manA, nagA, nagK*, and CHS7). This pool was diluted in PBS and 100μl was injected i.v. into mice. 100 μl was also directly plated on LB + kanamycin plates to serve as the input comparison. The bile, spleen, lungs, right kidney, and all four lobes of the liver (separately) were harvested and plated on 150mm LB+kanamycin plates. After overnight incubation at 37°, cells were scraped and processed as described above. Sequencing reads were mapped to the list of 21 barcodes and custom R scripts (available at https://github.com/hullahalli/stampr_rtisan) were used to quantify the change in barcode frequency relative to the input. Note that the method to generate barcoded libraries is the same as previously described (47, 49), although the analytical pipeline differs due to the necessity of visualizing clonal expansion.

### Statistical analysis

P-values reported in figures were obtained from GraphPad Prism 9. Figure legends indicate the statistical test used.

### Data and code availability

Barcode counts and TIS read counts, in addition to scripts required analyze these data, are available at https://github.com/hullahalli/stampr_rtisan

## Acknowledgements

This work was supported by an NSF Graduate Research Fellowship (K.H.) and NIH RO1 AI-042347 and Howard Hughes Medical Institute (M.K.W.). We are grateful to Bolutife Fakoya, Abdelrahim Zoued, Carole Kuehl, Ting Zhang, and Ian Campbell for assistance with plating organ homogenates. We are grateful to members of our lab and Michael Chao for providing feedback on the manuscript.

## Author Contributions

K.H. conceived, performed, and analyzed all experiments in the study. K.H. and M.K.W. designed experiments and wrote the manuscript.

## Figure Legends

<u>Figure S1. Barcode stability and creation of the STAMPR standard curve</u>

A) The barcoded library was serially passed in LB lacking antibiotic. Each day, colonies containing barcodes were enumerated as a fraction of the kanamycin-resistant cells. In addition, every day, 30 single colonies derived from the serially passaged culture were grown on plates lacking kanamycin and then patched onto media containing kanamycin. All colonies retained the antibiotic resistance marker, suggesting that the barcode is stable for at least ~50 generations. B) Growth curves of 9 individual barcoded clones (STAMP) and the WT untagged strain (CHS7) were indistinguishable, indicating that the tags do not confer a growth defect. C) Sequencing of the barcodes at different known bottleneck sizes (plated serial dilutions of the barcoded library) showed that both N_b_ and N_r_ accurately reflect FP values up to 10^5^ – corresponding to the approximate sequencing depth of the sample. The underlying barcode distributions that result from the serial dilutions are shown in D.

<u>Figure S2. Longitudinal CFU, N_b_, and N_r_ measurements from the blood, bile, and spleen</u>

Points with the same fill and border color are from the same animal; the same fill and border coloring scheme is used for the animals shown in Figure S3 and S4.

<u>Figure S3. Longitudinal CFU, N_b_, and N_r_ measurements from the lungs and kidneys</u>

Points with the same fill and border color are from the same animal and correspond to animals shown in Figure S2 and S4.

<u>Figure S4. Longitudinal CFU, N_b_, and N_r_ measurements from the liver</u>

Points with the same fill and border color are from the same animal and correspond to animals shown in Figure S2 and S3.

<u>Figure S5. Clonal expansion in the kidney.</u>

A) CFU of the right kidney is shown and is taken from the larger data set in Figure S3. Points with the same fill and border color are derived from the same animal; in animal B the right kidney possessed 10,000 times more bacteria than the left kidney, which was sterile. B) The underlying barcode distribution found in animal B. High CFU in the right kidney is attributable to marked expansion of 8 cells.

<u>Figure S6. GD and FRD 4 hpi (0 dpi)</u>

GD heatmaps are shown on the left and FRD heatmaps are shown on the right. Column names for FRD correspond to the reference sample. Organ codes are blood (a), bile (b) right spleen (e), left spleen (f), right kidney (j) left kidney (k), right lung (n), left lung (o), caudate liver (v), right liver (w), left liver (x), medial liver (y). In these animals, low GD values are reflective of their very high N_b_ values, as all samples are closely related to the inoculum. This is consistent with high FRD values.

<u>Figure S7. GD and FRD 1 dpi</u>

GD heatmaps are shown on the left and FRD heatmaps are shown on the right. Column names for FRD correspond to the reference sample. Organ codes are blood (a), bile (b) right spleen (e), left spleen (f), right kidney (j) left kidney (k), right lung (n), left lung (o), caudate liver (v), right liver (w), left liver (x), medial liver (y). By 1 dpi, populations in highly bottlenecked samples are largely unrelated to each other. The spleen and liver still possess modestly diverse populations and therefore have lower GD values. The “checkerboarding” effect in the liver samples (v,w,x,y: bottom right) of Mouse 5 and 7 are due to expanded clones in the liver that increase genetic distance across all samples (asterisks). The FRD metric was in part designed to detect these events, as comparisons with samples possessing clonal expansion events yielded both high GD and FRD, indicating the underlying population is similar and diverse. The single exception was observed in the right lobe of mouse 7. The clonal expansion in this sample was so profound that the underlying population is indistinguishable from noise (see Figure 2)

<u>Figure S8. GD and FRD 2 dpi</u>

GD heatmaps are shown on the left and FRD heatmaps are shown on the right. Column names for FRD correspond to the reference sample. Organ codes are blood (a), bile (b) right spleen (e), left spleen (f), right kidney (j) left kidney (k), right lung (n), left lung (o), caudate liver (v), right liver (w), left liver (x), medial liver (y). Most organs are now highly divergent (high GD), apart from the liver, where adjacent lobes share many tags and are diverse.

<u>Figure S9. GD and FRD 3 dpi</u>

GD heatmaps are shown on the left and FRD heatmaps are shown on the right. Column names for FRD correspond to the reference sample. Organ codes are blood (a), bile (b) right spleen (e), left spleen (f), right kidney (j) left kidney (k), right lung (n), left lung (o), caudate liver (v), right liver (w), left liver (x), medial liver (y). Relatedness between all organs has substantially decreased. Mouse 16 is the first animal where there is essentially no similarity between liver lobes, indicating that by 3 dpi clearance has completely segregated sub-organ populations. The checkerboarding in mouse 13 is due to transferred clones. Moderate FRD values indicate that transferred populations comprise multiple clones although underlying non-transferred populations are present as well.

<u>Figure S10. GD and FRD 4 dpi</u>

GD heatmaps are shown on the left and FRD heatmaps are shown on the right. Column names for FRD correspond to the reference sample. Organ codes are blood (a), bile (b) right spleen (e), left spleen (f), right kidney (j) left kidney (k), right lung (n), left lung (o), caudate liver (v), right liver (w), left liver (x), medial liver (y). A clear dichotomy is observed between Mice 18 and 19 versus mouse 17 and 20. Mice 17 and 20 had substantial replication of bacteria in the bile. This clone spread across the body, indicated by low GD values in multiple organs. FRD values reflect the fact that the bile is a single clone. Starting at the asterisk, rightward movement on the FRD heatmap yields lower nonzero FRD values than downward movement. This indicates that when the non-bile organ is set as the reference, FRD values are low, revealing that only a few barcodes are shared. When the bile is set as the reference, FRD values are much higher, indicating that the shared clones represent a larger fraction of barcodes in the bile. This is quantified in Figure 3C. Given that the N_r_ of bile is ~1, low GD values are due to essentially a single clone.

<u>Figure S11. GD and FRD for 5 dpi</u>

GD heatmaps are shown on the left and FRD heatmaps are shown on the right. Column names for FRD correspond to the reference sample. Organ codes are blood (a), bile (b) right spleen (e), left spleen (f), right kidney (j) left kidney (k), right lung (n), left lung (o), caudate liver (v), right liver (w), left liver (x), medial liver (y). Mice 23, 24, 32, 33, and 52 had abscesses in at least one lobe of the liver. Mouse 31 had a clone apparently disseminated from bile. Mouse 53 had much higher blood CFU than others in the cohort. Note that most instances of sharing correspond to low FRD values, indicating that sharing is due to very few barcodes. A notable exception is the splenic samples of mouse 23, which are represented in Figure 2.

<u>Figure S12. CFU and population structure from in vivo transposon mutant screen</u>

A) Bacterial burdens for the liver, spleen and bile are shown for animals sacrificed 1 dpi following inoculation of the transposon library. The three animals with bile CFU correspond to the three animals where spreading of clones between the liver and spleen was observed C,D. B-D) Frequencies of mutants (analogous to barcodes) are shown as a function of genome position. Mutants are relatively evenly represented in the inoculum (B). C and D) Distributions from the livers and spleens from five animals. 3 animals possessed marked expansion of a single mutant that spread between the liver and spleen (red triangles).

<u>Figure S13. CFU and additional organs from Figure 7.</u>

A) Bacterial burdens from the spleen, right kidney, lungs, and all four lobes of the liver are shown for 1 dpi and 5 dpi of the barcoded mutant library. High CFU in the liver 5dpi corresponded to animals with abscesses. Points with the same color and shape were obtained from the same animal. B) Fold change of barcode abundance of strains in the kidney are shown for 1 dpi (left) and 5 dpi right). Too few samples were obtained from 5 dpi kidney samples for analysis. C) Same as B) but for lobes of the liver 5 dpi; note this graph separates mutant fitness by liver lobe rather than presence or absence of liver abscesses, as done in Figure 7. P-values were obtained by Dunn’s multiple comparisons test.

<u>Figure S14. Schematic summary of key findings</u>

A and B schematize the most distinct trajectories of ExPEC population dynamics identified – one where bacteria are nearly cleared (A) and one where they expand clonally (B). Additional intermediate categories are also observed when clonally expanded populations disseminate systemically and alter bacterial burdens and population structure in distal organs (C). The probability of dissemination depends on the organ where clonal expansion occurred; e.g., there was a much larger likelihood of disseminating when there was clonal expansion evident in bile versus from the liver. Every animal displays a unique pattern of expansion and dissemination.

<u>Table S1. Genes identified by TIS</u>

<u>Table S2. Oligonucleotides</u>

<u>Table S3. Strains and plasmids</u>

## References

1. Christaki E, Giamarellos-Bourboulis EJ. 2014. The complex pathogenesis of bacteremia: from antimicrobial clearance mechanisms to the genetic background of the host. Virulence 5:57–65.

2. Jenne CN, Kubes P. 2013. Immune surveillance by the liver. Nat Immunol 14:996–1006.

3. Krenkel O, Tacke F. 2017. Liver macrophages in tissue homeostasis and disease. Nat Rev Immunol 17:306–321.

4. Smith SN, Hagan EC, Lane MC, Mobley HLT. 2010. Dissemination and systemic colonization of uropathogenic Escherichia coli in a murine model of bacteremia. MBio 1:e00262–10.

5. Armbruster CE, Forsyth VS, Johnson AO, Smith SN, White AN, Brauer AL, Learman BS, Zhao L, Wu W, Anderson MT, Bachman MA, Mobley HLT. 2019. Twin arginine translocation, ammonia incorporation, and polyamine biosynthesis are crucial for Proteus mirabilis fitness during bloodstream infection. PLoS Pathog 15:e1007653.

6. Holmes CL, Anderson MT, Mobley HLT, Bachman MA. 2021. Pathogenesis of Gram-Negative Bacteremia. Clin Microbiol Rev 34.

7. Ercoli G, Fernandes VE, Chung WY, Wanford JJ, Thomson S, Bayliss CD, Straatman K, Crocker PR, Dennison A, Martinez-Pomares L, Andrew PW, Moxon ER, Oggioni MR. 2018. Intracellular replication of Streptococcus pneumoniae inside splenic macrophages serves as a reservoir for septicaemia. Nat Microbiol 3:600–610.

8. Gresham HD, Lowrance JH, Caver TE, Wilson BS, Cheung AL, Lindberg FP. 2000. Survival of Staphylococcus aureus Inside Neutrophils Contributes to Infection. J Immunol 164:3713–3722.

9. Siggins MK, Lynskey NN, Lamb LE, Johnson LA, Huse KK, Pearson M, Banerji S, Turner CE, Woollard K, Jackson DG, Sriskandan S. 2020. Extracellular bacterial lymphatic metastasis drives Streptococcus pyogenes systemic infection. Nat Commun 11:1–12.

10. Dale AP, Woodford N. 2015. Extra-intestinal pathogenic Escherichia coli (ExPEC): Disease, carriage and clones. J Infect 71:615–626.

11. Abel S, Abel zur Wiesch P, Davis BM, Waldor MK. 2015. Analysis of Bottlenecks in Experimental Models of Infection. PLOS Pathog 11:e1004823.

12. Jorch SK, Surewaard BGJ, Hossain M, Peiseler M, Deppermann C, Deng J, Bogoslowski A, Wal F van der, Omri A, Hickey MJ, Kubes P. 2019. Peritoneal GATA6+ macrophages function as a portal for Staphylococcus aureus dissemination. J Clin Invest 129:4643–4656.

13. Fiebig A, Vrentas CE, Le T, Huebner M, Boggiatto PM, Olsen SC, Crosson S. 2020. Quantification of Brucella abortus population structure in a natural host. bioRxiv. bioRxiv.

14. Grant AJ, Restif O, McKinley TJ, Sheppard M, Maskell DJ, Mastroeni P. 2008. Modelling within-Host Spatiotemporal Dynamics of Invasive Bacterial Disease. PLoS Biol 6:e74.

15. Blundell JR, Levy SF. 2014. Beyond genome sequencing: Lineage tracking with barcodes to study the dynamics of evolution, infection, and cancer. Genomics 104:417–430.

16. Bachta KER, Allen JP, Cheung BH, Chiu C-H, Hauser AR. 2020. Systemic infection facilitates transmission of Pseudomonas aeruginosa in mice. Nat Commun 11:543.

17. Pollitt EJG, Szkuta PT, Burns N, Foster SJ. 2018. Staphylococcus aureus infection dynamics. PLOS Pathog 14:e1007112.

18. Gerlini A, Colomba L, Furi L, Braccini T, Manso AS, Pammolli A, Wang B, Vivi A, Tassini M, van Rooijen N, Pozzi G, Ricci S, Andrew PW, Koedel U, Moxon ER, Oggioni MR. 2014. The Role of Host and Microbial Factors in the Pathogenesis of Pneumococcal Bacteraemia Arising from a Single Bacterial Cell Bottleneck. PLoS Pathog 10:e1004026.

19. Jasinska W, Manhart M, Lerner J, Gauthier L, Serohijos AWR, Bershtein S. 2020. Chromosomal barcoding of E. coli populations reveals lineage diversity dynamics at high resolution. Nat Ecol Evol 4:437–452.

20. Abel S, Abel zur Wiesch P, Chang H-H, Davis BM, Lipsitch M, Waldor MK. 2015. Sequence tag–based analysis of microbial population dynamics. Nat Methods 12:223–226.

21. Mahmutovic A, Gillman AN, Lauksund S, Robson Moe NA, Manzi A, Storflor M, Abel zur Wiesch P, Abel S. 2021. RESTAMP – Rate estimates by sequence-tag analysis of microbial populations. Comput Struct Biotechnol J 19:1035–1051.

22. Zhang T, Sasabe J, Hullahalli K, Sit B, Waldor MK. 2021. Increased Listeria monocytogenes dissemination and altered population dynamics in Muc2-deficient mice. Infect Immun https://doi.org/10.1128/IAI.00667-20.

23. Zhang T, Abel S, Abel Zur Wiesch P, Sasabe J, Davis BM, Higgins DE, Waldor MK. 2017. Deciphering the landscape of host barriers to Listeria monocytogenes infection. Proc Natl Acad Sci U S A 114:6334–6339.

24. Liu X, Kimmey JM, Matarazzo L, de Bakker V, Van Maele L, Sirard JC, Nizet V, Veening JW. 2021. Exploration of Bacterial Bottlenecks and Streptococcus pneumoniae Pathogenesis by CRISPRi-Seq. Cell Host Microbe 29:107–120.e6.

25. Hullahalli K, Pritchard JR, Waldor MK. 2021. Refined quantification of infection bottlenecks and pathogen dissemination with STAMPR. bioRxiv 2021.04.28.441820.

26. Diabate M, Munro P, Garcia E, Jacquel A, Michel G, Obba S, Goncalves D, Luci C, Marchetti S, Demon D, Degos C, Bechah Y, Mege J-L, Lamkanfi M, Auberger P, Gorvel J-P, Stuart LM, Landraud L, Lemichez E, Boyer L. 2015. Escherichia coli α-Hemolysin Counteracts the Anti-Virulence Innate Immune Response Triggered by the Rho GTPase Activating Toxin CNF1 during Bacteremia. PLOS Pathog 11:e1004732.

27. McAdow M, Kim HK, DeDent AC, Hendrickx APA, Schneewind O, Missiakas DM. 2011. Preventing Staphylococcus aureus Sepsis through the Inhibition of Its Agglutination in Blood. PLoS Pathog 7:e1002307.

28. Krimbas CB, Tsakas S. 1971. THE GENETICS OF *DACUS OLEAE*. V. CHANGES OF ESTERASE POLYMORPHISM IN A NATURAL POPULATION FOLLOWING INSECTICIDE CONTROL-SELECTION OR DRIFT? Evolution (N Y) 25:454–460.

29. Welch RA, Burland V, Plunkett G, Redford P, Roesch P, Rasko D, Buckles EL, Liou S-R, Boutin A, Hackett J, Stroud D, Mayhew GF, Rose DJ, Zhou S, Schwartz DC, Perna NT, Mobley HLT, Donnenberg MS, Blattner FR. 2002. Extensive mosaic structure revealed by the complete genome sequence of uropathogenic Escherichia coli. Proc Natl Acad Sci U S A 99:17020–4.

30. Ristow LC, Welch RA. 2016. Hemolysin of uropathogenic Escherichia coli: A cloak or a dagger? Biochim Biophys Acta - Biomembr 1858:538–545.

31. Hryckowian AJ, Welch RA. 2013. RpoS contributes to phagocyte oxidase-mediated stress resistance during urinary tract infection by Escherichia coli CFT073. MBio 4.

32. Marcus S, Esplin DW, Donaldson DM. 1954. Lack of bactericidal effect of mouse serum on a number of common microorganisms. Science (80-) 119:877.

33. Sanchez KK, Chen GY, Schieber AMP, Redford SE, Shokhirev MN, Leblanc M, Lee YM, Ayres JS. 2018. Cooperative Metabolic Adaptations in the Host Can Favor Asymptomatic Infection and Select for Attenuated Virulence in an Enteric Pathogen. Cell 175:146–158.e15.

34. Schmid-Hempel P, Frank SA. 2007. Pathogenesis, Virulence, and Infective Dose. PLoS Pathog 3:e147.

35. Llorente C, Schnabl B. 2016. Fast-Track Clearance of Bacteria from the Liver. Cell Host Microbe 20:1–2.

36. Rooijen N Van, Sanders A. 1994. Liposome mediated depletion of macrophages: mechanism of action, preparation of liposomes and applications. J Immunol Methods 174:83–93.

37. Tavares AJ, Poon W, Zhang YN, Dai Q, Besla R, Ding D, Ouyang B, Li A, Chen J, Zheng G, Robbins C, Chan WCW, Murphy CJ. 2017. Effect of removing Kupffer cells on nanoparticle tumor delivery. Proc Natl Acad Sci U S A 114:E10871–E10880.

38. Malinverni JC, Silhavy TJ. 2009. An ABC transport system that maintains lipid asymmetry in the gram-negative outer membrane. Proc Natl Acad Sci U S A 106:8009–14.

39. Ekiert DC, Bhabha G, Isom GL, Greenan G, Ovchinnikov S, Henderson IR, Cox JS, Vale RD. 2017. Architectures of Lipid Transport Systems for the Bacterial Outer Membrane. Cell 169:273–285.e17.

40. Du S, Pichoff S, Lutkenhaus J. 2016. FtsEX acts on FtsA to regulate divisome assembly and activity. Proc Natl Acad Sci U S A 113:E5052–E5061.

41. Yang DC, Peters NT, Parzych KR, Uehara T, Markovski M, Bernhardt TG. 2011. An ATP-binding cassette transporter-like complex governs cell-wall hydrolysis at the bacterial cytokinetic ring. Proc Natl Acad Sci U S A 108:E1052–E1060.

42. Hall MN, Silhavy TJ. 1981. Genetic analysis of the ompB locus in Escherichia coli K-12. J Mol Biol 151:1–15.

43. Paixão L, Oliveira J, Veríssimo A, Vinga S, Lourenço EC, Ventura MR, Kjos M, Veening JW, Fernandes VE, Andrew PW, Yesilkaya H, Neves AR. 2015. Host glycan sugar-specific pathways in streptococcus pneumonia: Galactose as a key sugar in colonisation and infection. PLoS One 10.

44. Uehara T, Park JT. 2004. The N-acetyl-D-glucosamine kinase of Escherichia coli and its role in murein recycling. J Bacteriol 186:7273–7279.

45. Karlinsey JE, Stepien TA, Mayho M, Singletary LA, Bingham-Ramos LK, Brehm MA, Greiner DL, Shultz LD, Gallagher LA, Bawn M, Kingsley RA, Libby SJ, Fang FC. 2019. Genome-wide Analysis of Salmonella enterica serovar Typhi in Humanized Mice Reveals Key Virulence Features. Cell Host Microbe 26:426–434.e6.

46. Wang N, Ozer EA, Mandel MJ, Hauser AR. 2014. Genome-wide identification of Acinetobacter baumannii genes necessary for persistence in the lung. MBio 5:1163–1177.

47. Warr AR, Hubbard TP, Munera D, Blondel CJ, Abel zur Wiesch P, Abel S, Wang X, Davis BM, Waldor MK. 2019. Transposon-insertion sequencing screens unveil requirements for EHEC growth and intestinal colonization. PLOS Pathog 15:e1007652.

48. Subashchandrabose S, Smith SN, Spurbeck RR, Kole MM, Mobley HLT. 2013. Genome-Wide Detection of Fitness Genes in Uropathogenic Escherichia coli during Systemic Infection. PLoS Pathog 9:e1003788.

49. Hubbard TP, Chao MC, Abel S, Blondel CJ, Wiesch PA Zur, Zhou X, Davis BM, Waldor MK. 2016. Genetic analysis of vibrio parahaemolyticus intestinal colonization. Proc Natl Acad Sci U S A 113:6283–6288.

50. Xu SX, Gilmore KJ, Szabo PA, Zeppa JJ, Baroja ML, Haeryfar SMM, McCormick JK. 2014. Superantigens subvert the neutrophil response to promote abscess formation and enhance Staphylococcus aureus survival in vivo. Infect Immun 82:3588–3598.

51. Cheng AG, Hwan KK, Burts ML, Krausz T, Schneewind O, Missiakas DM. 2009. Genetic requirements for Staphylococcus aureus abscess formation and persistence in host tissues. FASEB J 23:3393–3404.

52. Subashchandrabose S, Smith S, DeOrnellas V, Crepin S, Kole M, Zahdeh C, Mobley HLT. 2016. Acinetobacter baumannii Genes Required for Bacterial Survival during Bloodstream Infection. mSphere 1:e00013–15.

53. Anderson MT, Mitchell LA, Zhao L, Mobley HLT. 2018. Citrobacter freundii fitness during bloodstream infection. Sci Rep 8:11792.

54. Anderson MT, Mitchell LA, Zhao L, Mobley HLT. 2017. Capsule Production and Glucose Metabolism Dictate Fitness during Serratia marcescens Bacteremia. MBio 8:e00740–17.

55. McCarthy AJ, Stabler RA, Taylor PW. 2018. Genome-wide identification by transposon insertion sequencing of Escherichia coli K1 genes essential for in vitro growth, gastrointestinal colonizing capacity, and survival in serum. J Bacteriol 200.

56. Cunrath O, Bumann D. 2019. Host resistance factor SLC11A1 restricts Salmonella growth through magnesium deprivation. Science (80-) 366:995–999.

57. Ellner SP, Buchon N, Orr TD, Lazzaro BP. 2021. Host-pathogen Immune Feedbacks Can Explain Widely Divergent Outcomes from Similar Infections. bioRxiv 2021.01.08.425954.

58. Duneau D, Ferdy JB, Revah J, Kondolf H, Ortiz GA, Lazzaro BP, Buchon N. 2017. Stochastic variation in the initial phase of bacterial infection predicts the probability of survival in D. melanogaster. Elife 6.

59. Crépin S, Ottosen EN, Peters K, Smith SN, Himpsl SD, Vollmer W, Mobley HLT. 2018. The lytic transglycosylase MltB connects membrane homeostasis and in vivo fitness of Acinetobacter baumannii. Mol Microbiol 109:745–762.

60. Hryckowian AJ, Baisa GA, Schwartz KJ, Welch RA. 2015. DsdA does not affect colonization of the murine urinary tract by Escherichia coli CFT073. PLoS One 10.

61. Ferrières L, Hémery G, Nham T, Guérout AM, Mazel D, Beloin C, Ghigo JM. 2010. Silent mischief: Bacteriophage Mu insertions contaminate products of Escherichia coli random mutagenesis performed using suicidal transposon delivery plasmids mobilized by broad-host-range RP4 conjugative machinery. J Bacteriol 192:6418–6427.

62. Philippe N, Alcaraz JP, Coursange E, Geiselmann J, Schneider D. 2004. Improvement of pCVD442, a suicide plasmid for gene allele exchange in bacteria. Plasmid 51:246–255.

63. Jiang Y, Chen B, Duan C, Sun B, Yang J, Yang S. 2015. Multigene editing in the Escherichia coli genome via the CRISPR-Cas9 system. Appl Environ Microbiol 81:2506–14.

64. Donnenberg MS, Kaper JB. 1991. Construction of an eae deletion mutant of enteropathogenic Escherichia coli by using a positive-selection suicide vector. Infect Immun 59:4310–4317.

65. Chiang SL, Rubin EJ. 2002. Construction of a mariner-based transposon for epitope-tagging and genomic targeting. Gene 296:179–185.

66. Pritchard JR, Chao MC, Abel S, Davis BM, Baranowski C, Zhang YJ, Rubin EJ, Waldor MK. 2014. ARTIST: High-Resolution Genome-Wide Assessment of Fitness Using Transposon-Insertion Sequencing. PLoS Genet 10:e1004782.

67. Hubbard TP, Billings G, Dörr T, Sit B, Warr AR, Kuehl CJ, Kim M, Delgado F, Mekalanos JJ, Lewnard JA, Waldor MK. 2018. A live vaccine rapidly protects against cholera in an infant rabbit model. Sci Transl Med 10:eaap8423.

68. Hubbard TP, D’Gama JD, Billings G, Davis BM, Waldor MK. 2019. Unsupervised Learning Approach for Comparing Multiple Transposon Insertion Sequencing Studies. mSphere 4.

69. Subramaniyam S, Dejesus MA, Zaveri A, Smith CM, Baker RE, Ehrt S, Schnappinger D, Sassetti CM, Ioerger TR. 2019. Statistical analysis of variability in TnSeq data across conditions using zero-inflated negative binomial regression. BMC Bioinformatics 20:603.

