## Supplementary figures and images for "Pathogen clonal expansion underlies multiorgan dissemination and organ-specific outcomes during systemic infection"

### Figure S1

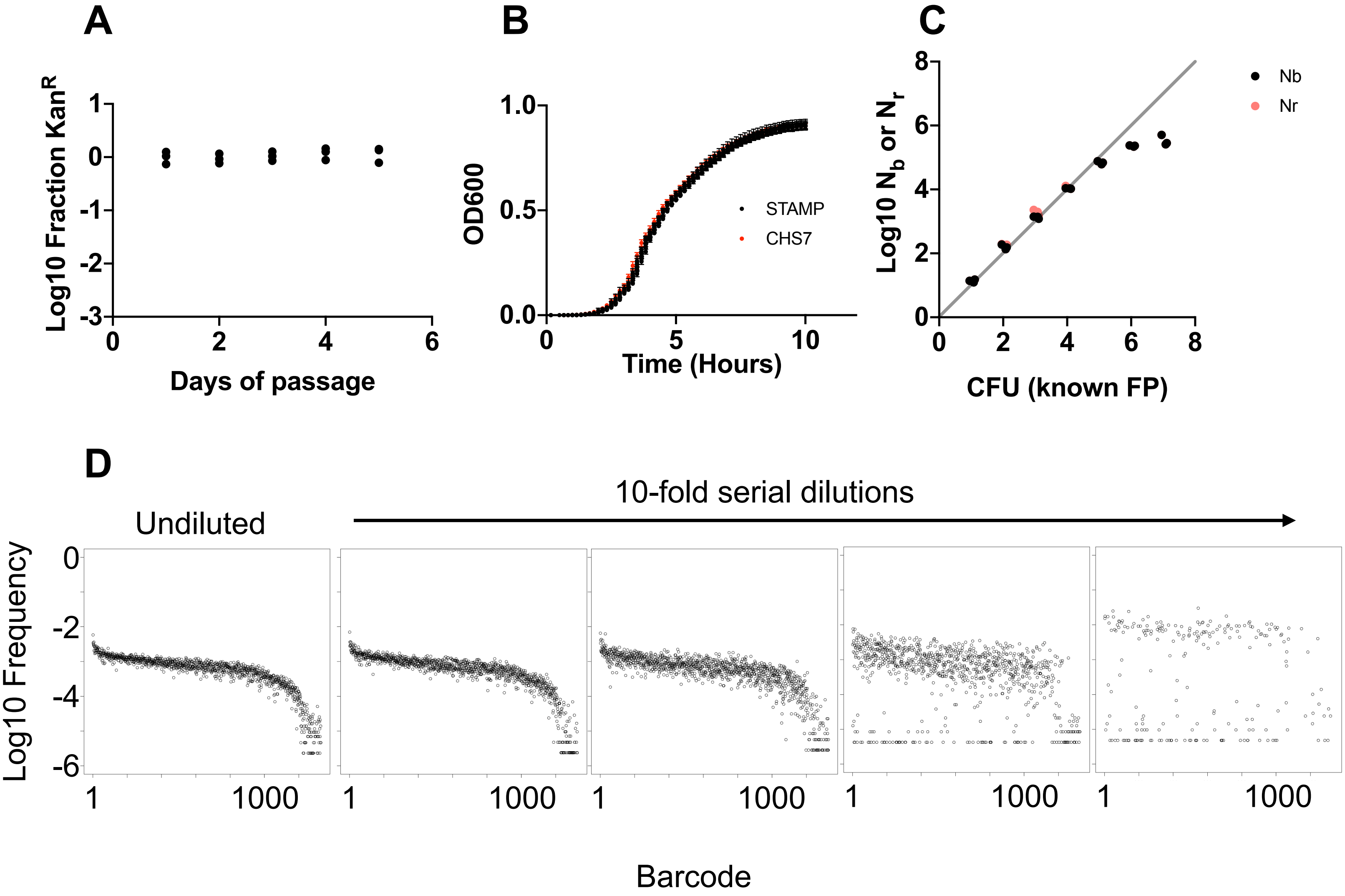

### Figure S2

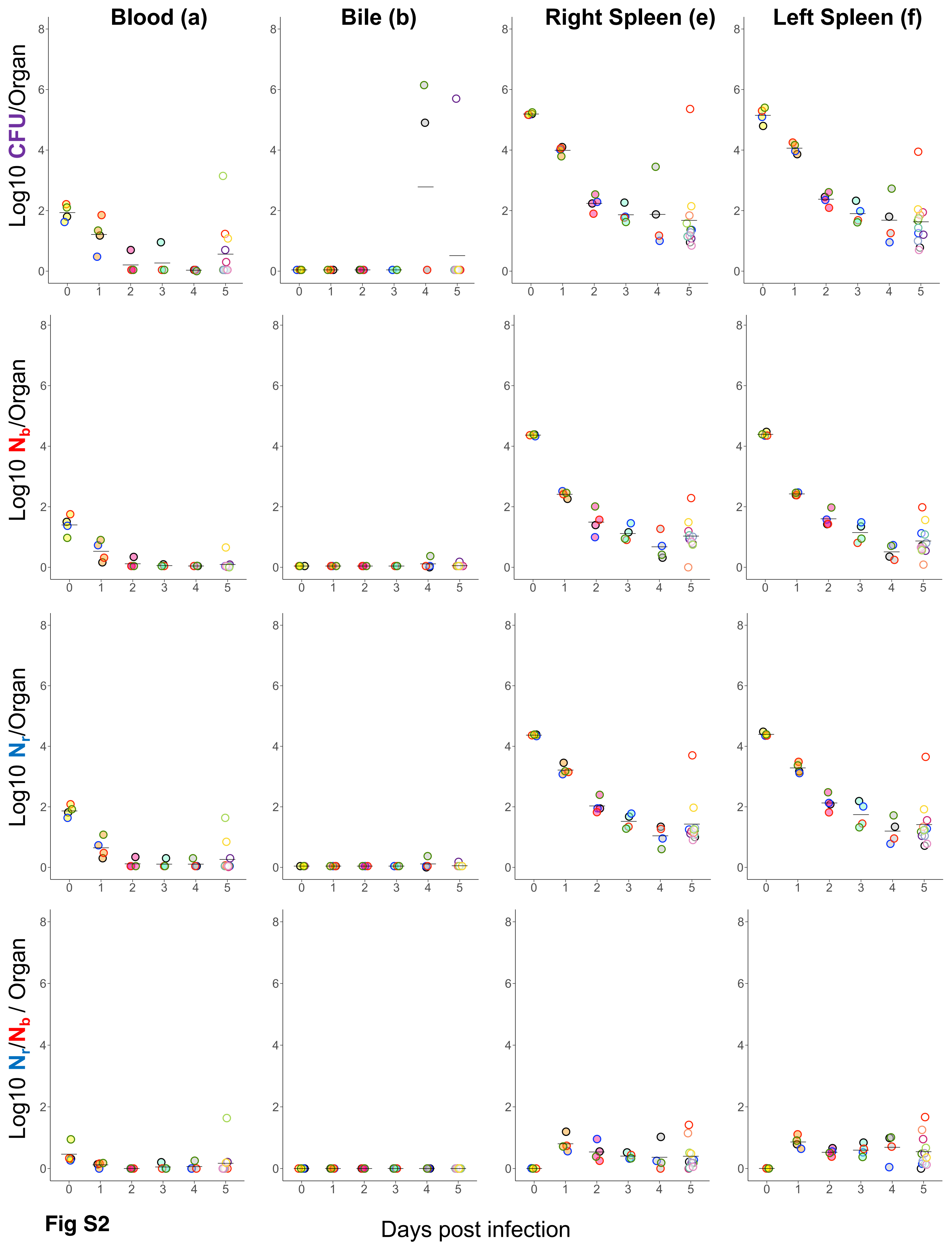

### Figure S3

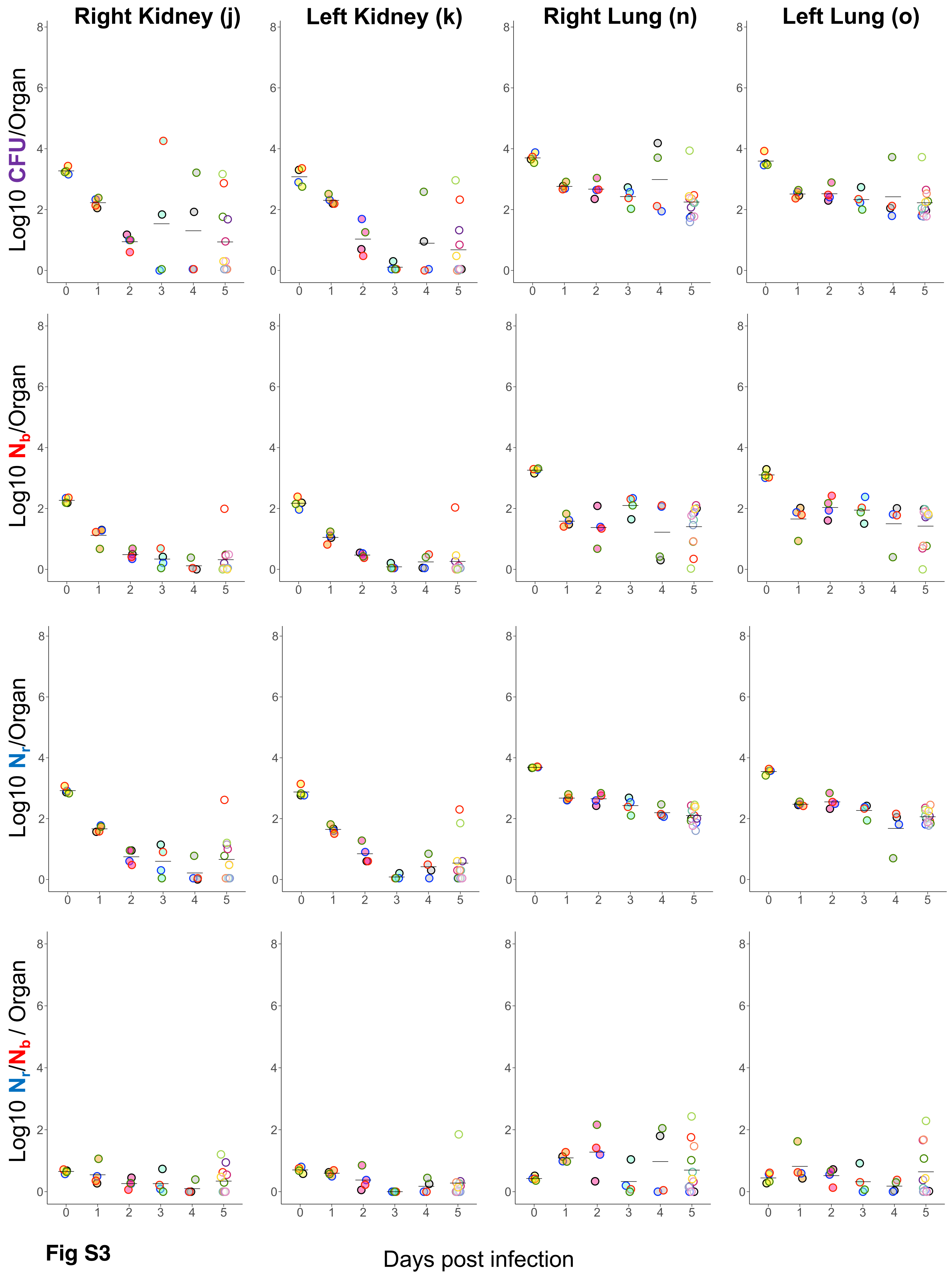

### Figure S4

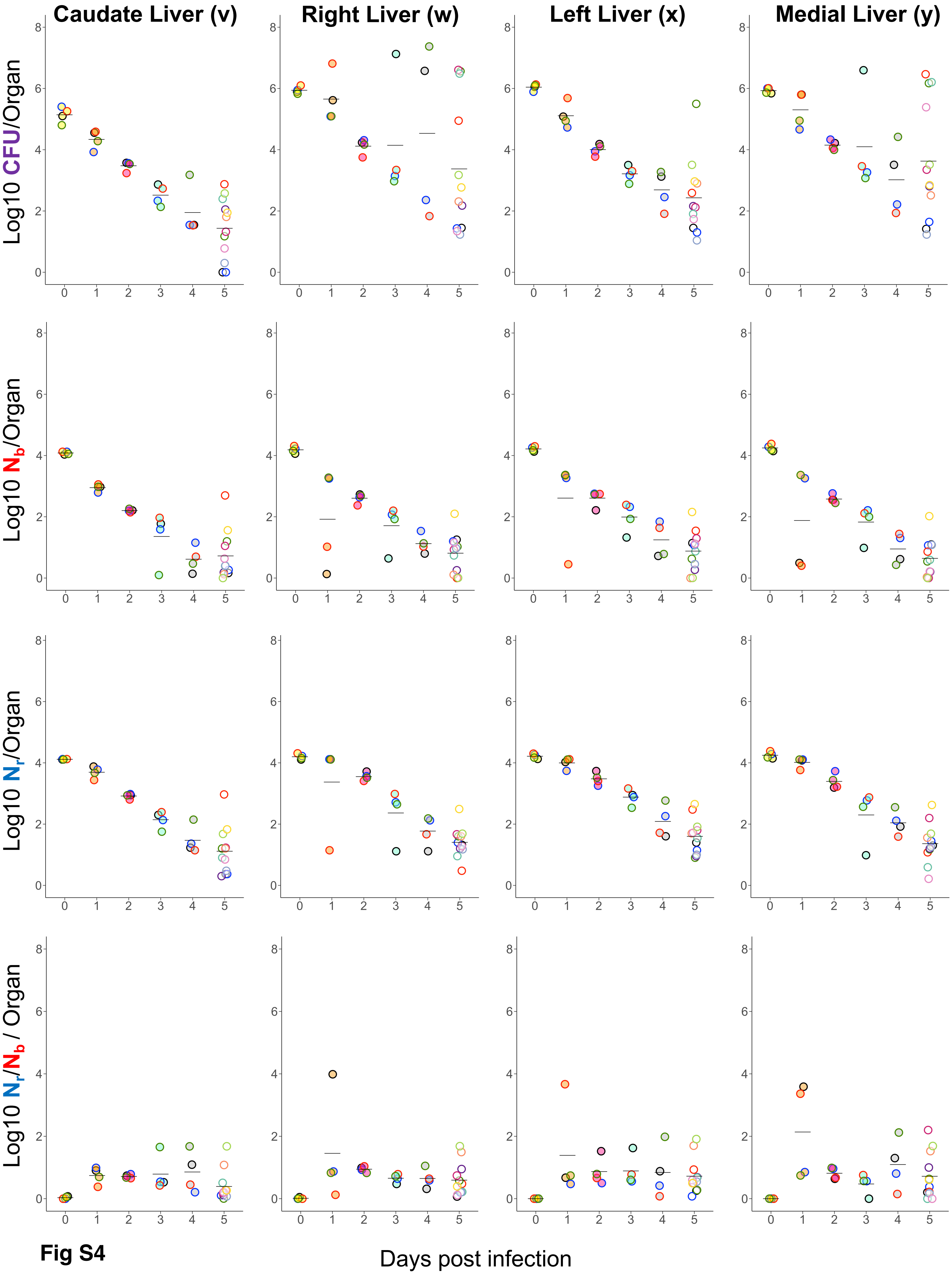

### Figure S5

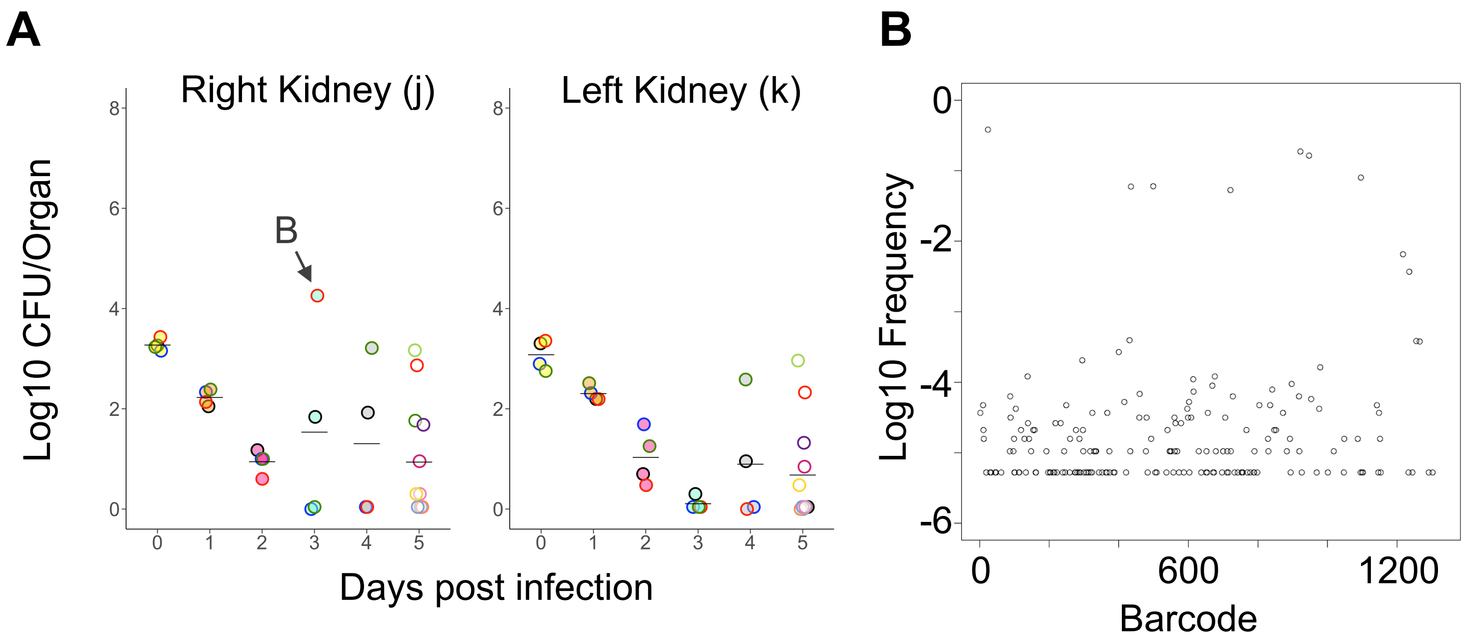

### Figure S11

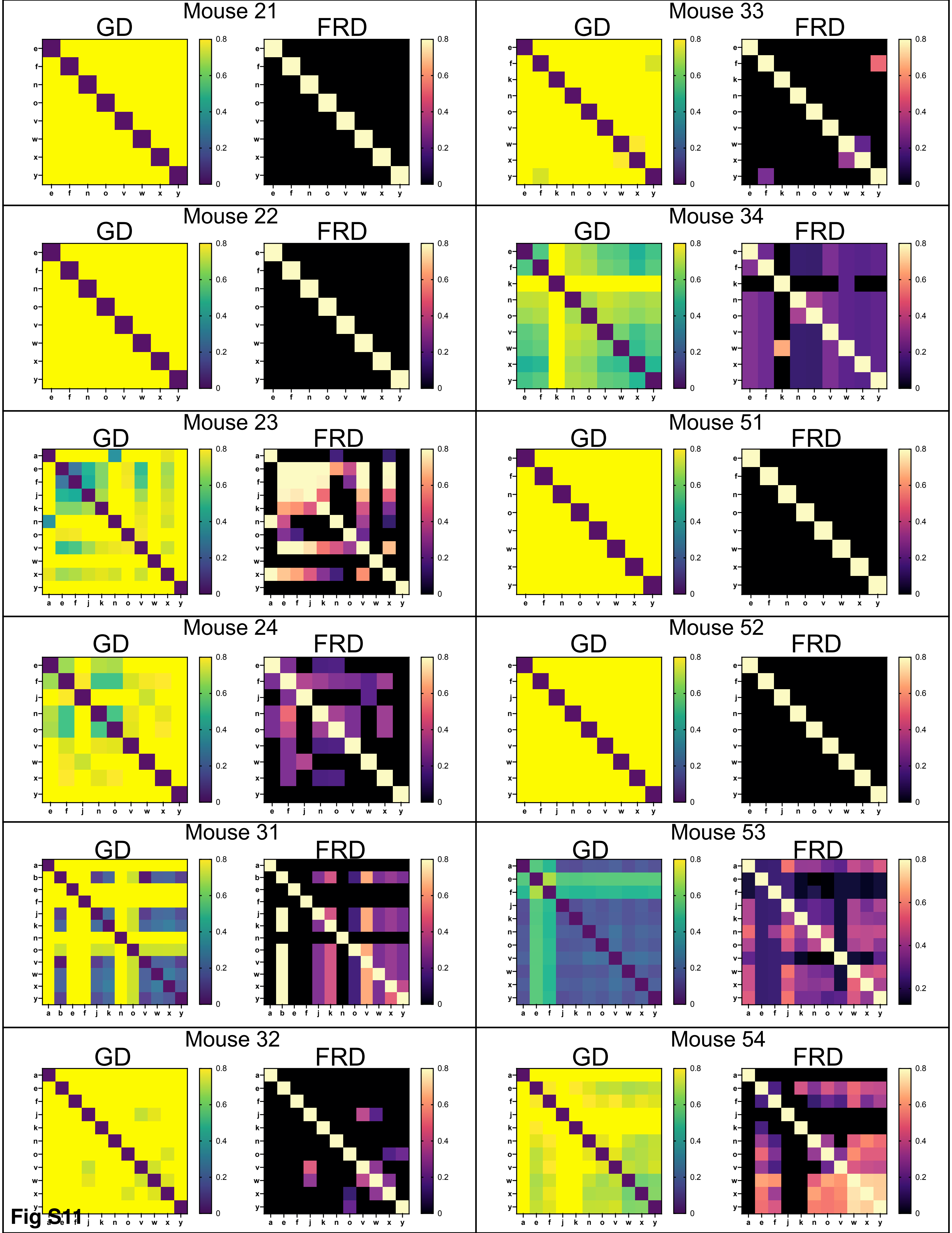

### Figure S12

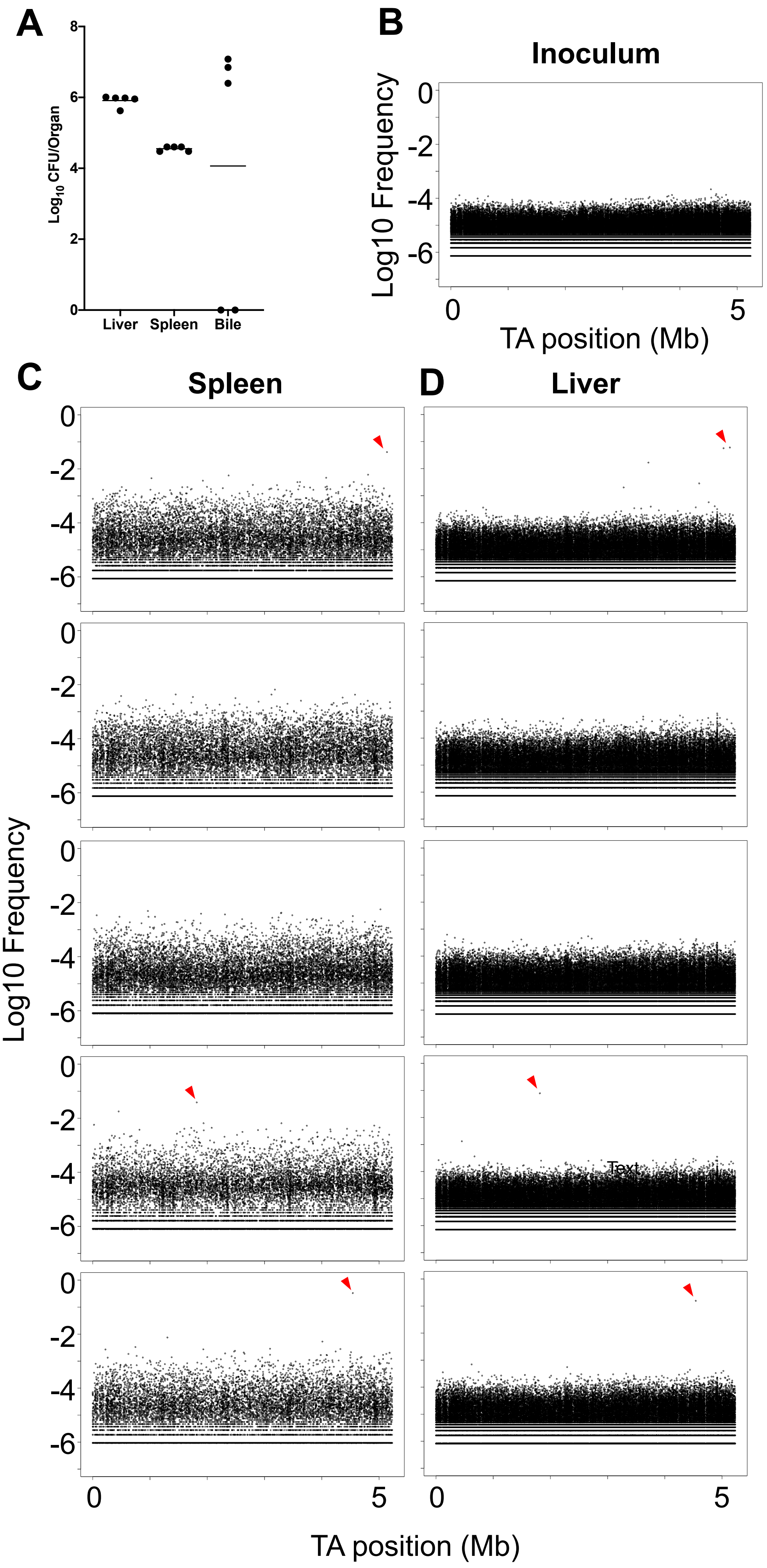

### Figure S13

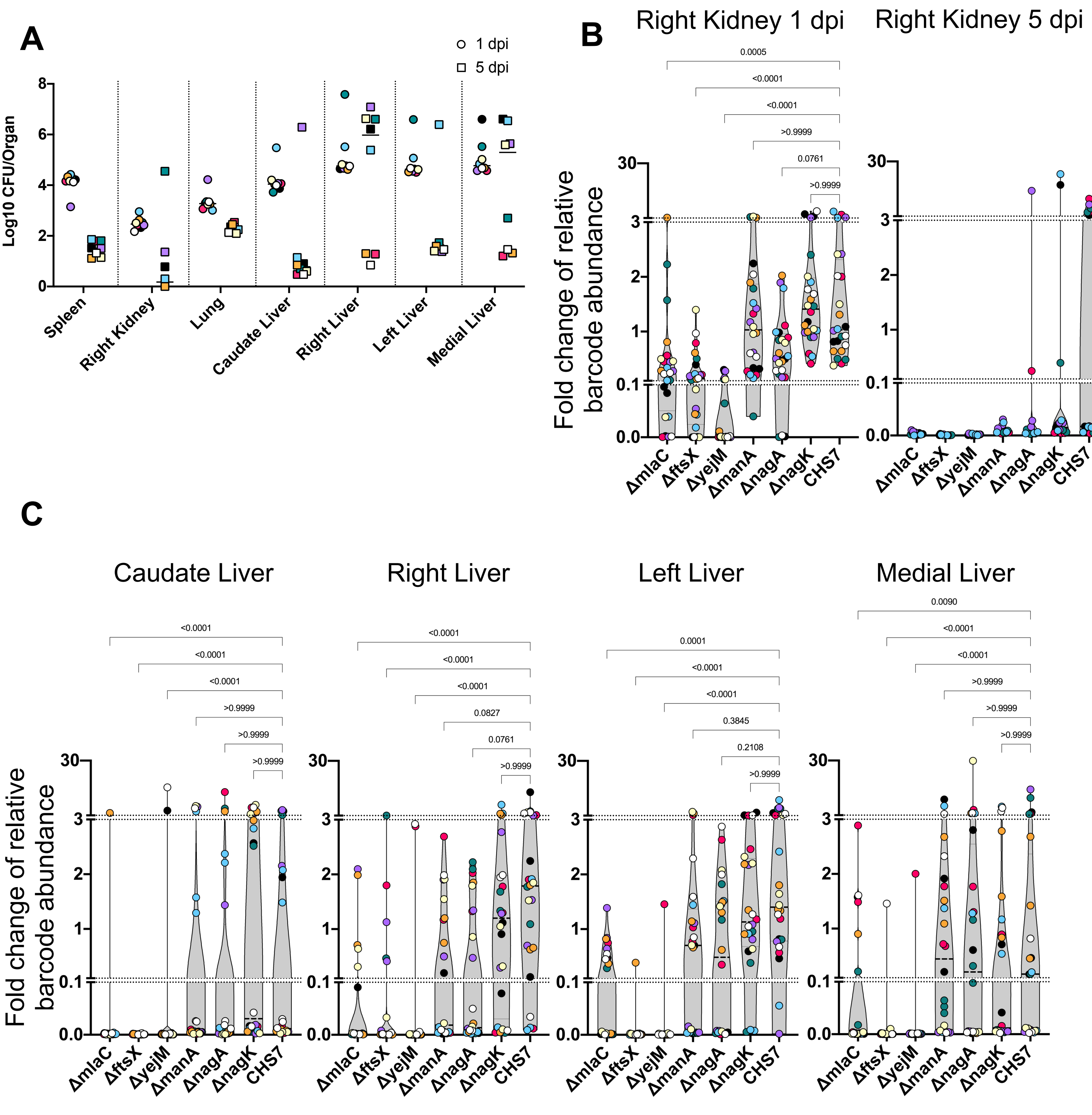

### Figure S14

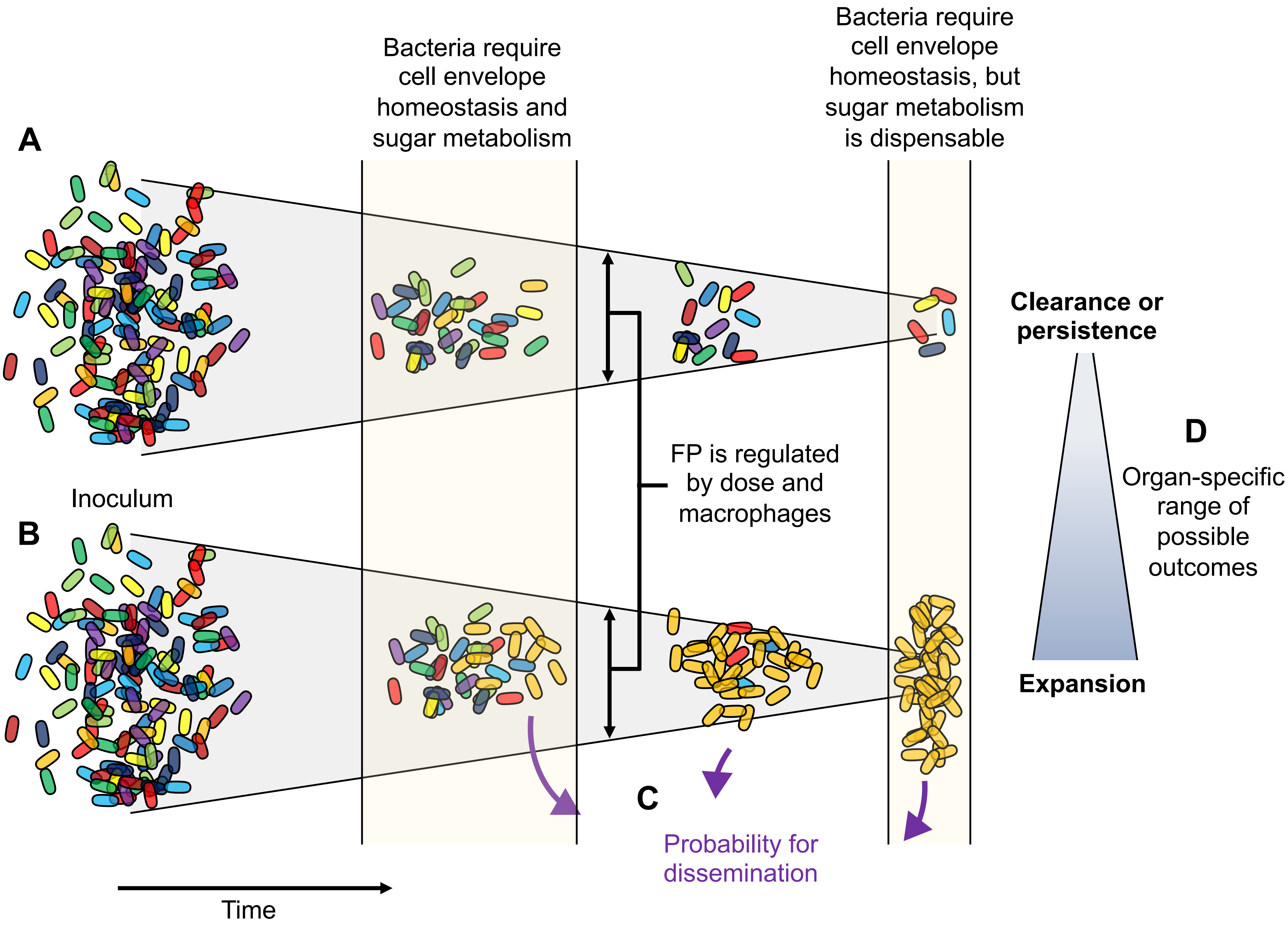
