## Supplementary material for "Pathogen clonal expansion underlies multiorgan dissemination and organ-specific outcomes during systemic infection": Figure S9

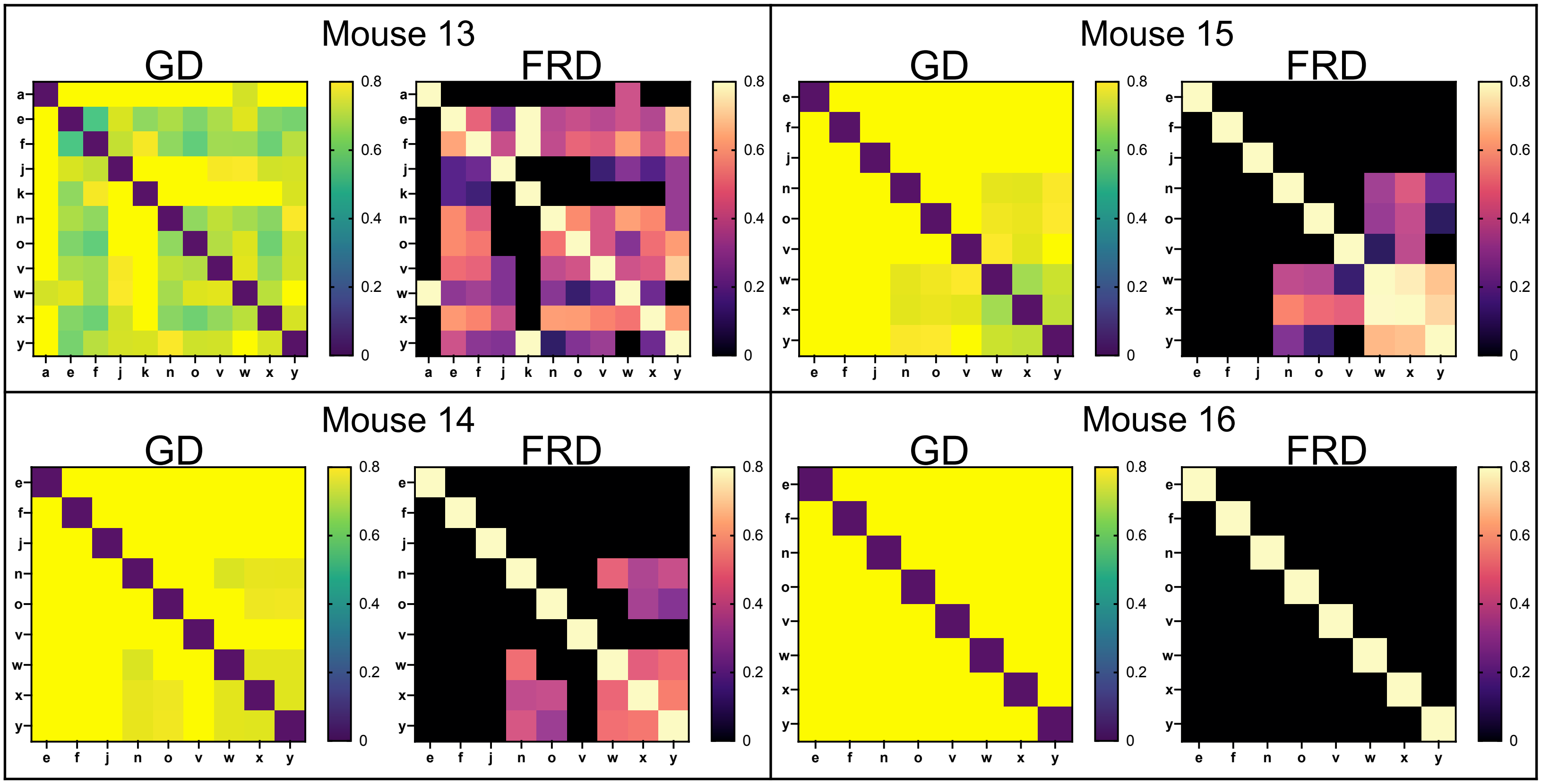


**Codes**

a - Blood (cardiac puncture)

b - Bile (aspirated from gallbladder)

e - Spleen (right half)

f - Spleen (left half)

j - Kidney (right)

k - Kidney (left)

n - Lung (right)

o - Lung (left)

v - Liver (caudate lobe)

w - Liver (right lobe)

x - Liver (left lobe)

y - Liver (medial lobe)

**Fig S9**
